# Prepatterned Tissue Stiffness Gradient Controls Organ shape and size

**DOI:** 10.64898/2026.09.04.748946

**Authors:** Jie Tong, Meng-Ju Lin, Nanzhong Deng, Marisa Delliturri, Mingang Xu, Sarah Millar, Mayumi Ito, Haogang Cai, Catherine P. Lu

## Abstract

Organ morphogenesis is orchestrated by precise coupling of mechanical and biochemical signals, yet how these two signals are integrated within tissue niche to regulate organ shape and size remains poorly understood. Here, using sweat gland as a tractable model of exocrine mini-organs, we demonstrate that dermal fibroblasts play a key role to translate prepatterned tissue stiffness gradients into spatial Wnt5a gradients via Piezo1-mediated mechanosensing. Following epidermal placode formation, progenitor cells collectively invaginate into the dermis, where spatiotemporally graded Wnt5a directs their sequential transition from ductal to glandular fate in softer distal dermis, specifying distinct exocrine compartments. Perturbation of dermal stiffness gradients and mechanosensing *in vivo* and *ex vivo* alters Wnt5a expression levels and gradient patterns and thereby changes the ductal length and glandular size, revealing a direct link between tissue mechanics and organ dimensions. To recapitulate this process *in vitro*, we engineer a microfluidic platform capable of generating robust and linear morphogen gradients in 3D, allowing duct-to-gland fate transitions to occur at a high resolution. Together, our finding identifies prepatterned stiffness gradients as a master upstream regulator of coupled mechanical-biochemical signaling, define a mechanistic framework for two-step sequential glandular morphogenesis, and offer new strategies for engineering glandular epithelia in regenerative medicine.

## Introduction

Organ architecture and sequential morphogenesis are not directly encoded in the genome but instead emerges through the precise coupling of mechanical cues with biochemical signals^1,2^. Mechanical force act as a central regulators of stem cells behavior – controlling positioning, shape, and fate – through translating mechanical cues into biochemical signals until structural equilibrium is achieved^3,4^. While considerable progress has been made in understanding how forces govern cell-cell or cell-ECM interactions, cytoskeletal rearrangement in organoids and cultured stem cells^5–8^, investigating how mechanical inputs shape organ architecture at different developmental stages *in vivo* remains challenging.

The intestinal villus has served as a robust model for studying morphogenesis driven by mechanical force^9,10^. Villus formation is initiated by compressive buckling forces from underlying endoderm, which patterns epithelial folding and influences morphogen gradient to establish in the stem cell niche^11,12^. More recently, active dewetting forces from mesenchyme have also been shown to contribute to this process^13^. Although morphogen gradients and a stiffness gradient along the crypt-villus axis have been independently proposed and partially measured^14,15^, whether and how these gradients are coordinated within the niche to specify stem cell fate during morphogenesis are still unclear.

Instead of folding and giving rise to various cell types within a single-layered epithelium, skin epithelium stratifies and forms various appendages (mini-organs) including tooth, hairs, sweat and mammary glands with distinct functions. They are initiated through evolutionarily conserved epithelial–mesenchymal interactions at epidermal placodes^16^. Early placode formation is driven by Wnt and Eda signaling from the epidermis^17–20^, while subsequent antagonism between epidermis-derived Shh and dermis-derived Bmp specifies cell fate toward distinct appendage types^21,22^. The mechanical contribution of these events has only recently been appreciated: contractile force across the epithelial–mesenchymal interface contributes to placode cell fate and downgrowth^23^; opposing dermal Fgf and Bmp signaling creates differential mechanical states that drive dermal condensation, which in turn mechanically activates β-catenin in overlying epidermal progenitors to initiate placode formation^24–26^. Yet, the coordination of mechanical and biochemical signals *beyond* the placode stage to drive 3D morphogenesis *in vivo* has not been established.

Using sweat gland as a tractable model of exocrine mini-organ morphogenesis, we demonstrated that a prepatterned stiffness gradient is converted into a morphogen gradient by fibroblasts through Piezo1-mediated mechanosensing. Beyond fate specification at the placode stage, the ductal progenitors collectively invaginate into dermis and experience a spatiotemporally decreased morphogen concentration, which instructs their fate transition from ductal to glandular identity. Fine-tuning of the stiffness gradient alters ductal length and glandular size, revealing a direct link between tissue mechanics and organ dimensions. Strikingly, this mechanism is shared by mammary, lacrimal, and salivary glands, suggesting a unifying principle whereby spatiotemporal coordination of mechanical and biochemical gradients controls exocrine gland morphogenesis — and potentially organ morphogenesis more broadly.

## Results

### Glandular morphology changes in response to dermal stiffness

To understand whether mechanical properties of dermal environment may impact on glandular morphogenesis, we used atomic force microscopy (AFM) (Figure S1A) to measure tissue stiffness in various glands at postnatal day 6 (P6), including sweat gland, lacrimal gland, mammary gland and salivary gland. We found that both sweat and lacrimal glands exhibit a stiffness gradient, highest in the ductal region and progressively decreasing toward the glandular portion (Figures 1A-B). In contrast, mammary and salivary glands, featuring short ducts and enlarged glandular structures, exhibited uniformly low stiffness compared to sweat and lacrimal glands (Figures 1C and S1B). To investigate the mechanisms governing ductal length and glandular size, we leveraged the sweat gland as a model system, whose developmental trajectory in mouse footpads is precisely staged postnatally -- progressing from placode formation through straight duct elongation (P0-P3) to coiled gland morphogenesis (P3-P6) (Figure S1C)^27^. To determine whether the stiffness gradient is an intrinsic property of the mesenchyme rather than a gland-induced phenomenon, we performed AFM measurements in regions devoid of glands. Strikingly, the upper dermis exhibited higher stiffness that declined progressively with depth, recapitulating the gradient observed in gland-containing areas (Figure S1D). Together, these results indicate the stiffness gradient as a pre-patterned feature of the mesenchyme, established prior to glandular morphogenesis.

**Figure 1.**
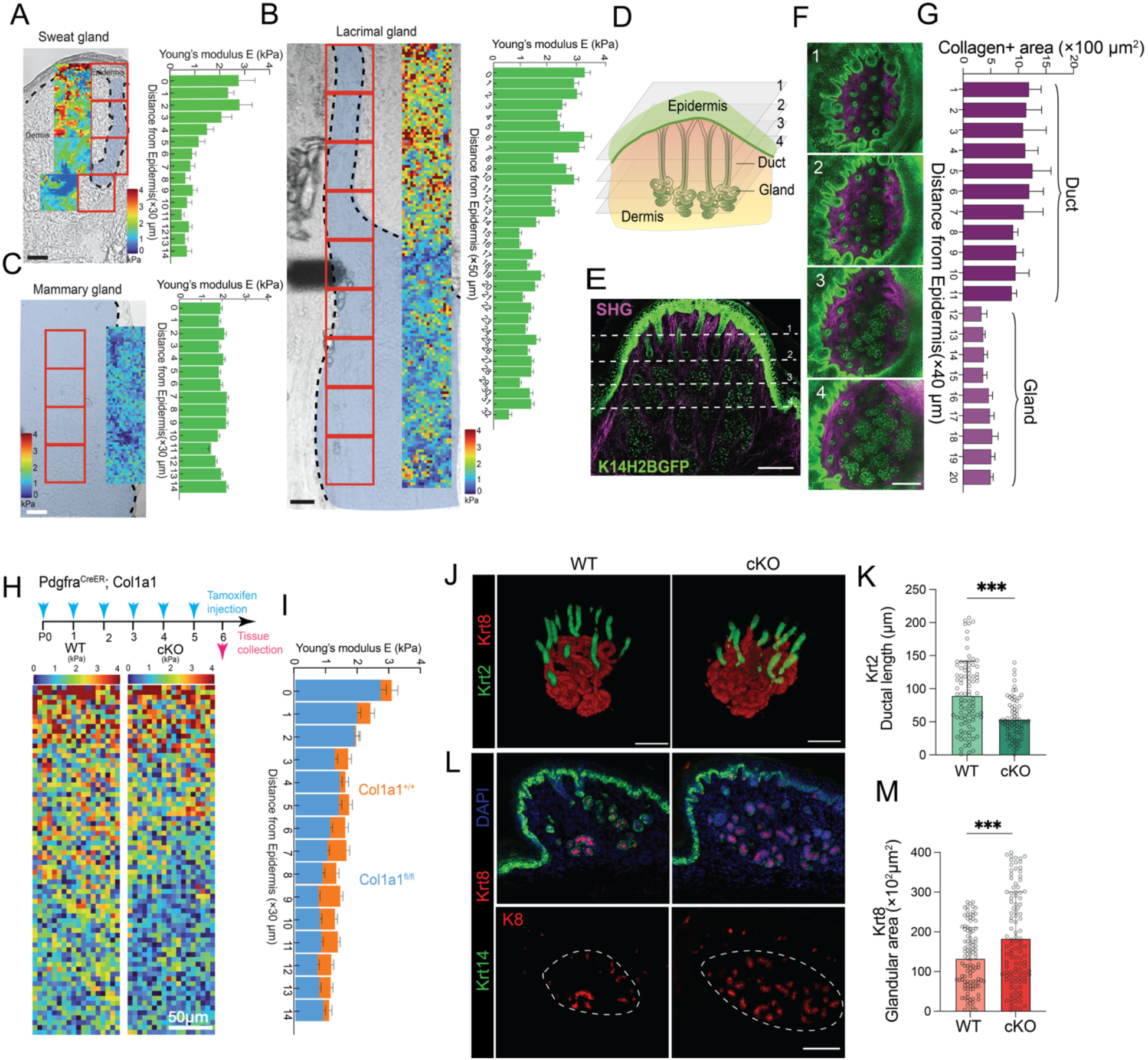
Sweat gland morphology changes in response to dermal stiffness. A-C. Spatial distribution of Young’s modulus E from the epidermis on the top to the deeper glandular areas in P6 mouse A) sweat gland; B) lacrimal gland; C) mammary gland, visualized as color-coded heat maps and bar graphs, showing the quantified values in kPa (Pascals) and aligned according to their corresponding distances from epidermis. Blue shading denotes the exocrine gland region, and red boxes indicate the areas measured. n = 4 mice. Scale bar, 50 µm. D. Schematic illustrating the process of whole-mount SHG imaging of P6 mouse foot skin, with Z-stack frames (1-4) acquired perpendicular to the direction of epidermis-dermis axis, corresponding to 1-4 in E and F. E-F. Fluorescent images from whole-mount SHG imaging, longitudinal and transverse sections, respectively. Scale bar, 100 µm. G. Quantification of collagen-positive (collagen+) area within 10 μm from the basement membrane, aligned according to their distances from the epidermis. Collagen-positive area was quantified within a 20 µm region surrounding the duct or gland epithelium. n = 3 mice. H-I. Pdgfra^CreER^;Col1a1^+/+^ (WT, n = 3 mice) and Pdgfra^CreER^;Col1a1^fl/fl^ (cKO, n = 3 mice) mice were induced with tamoxifen from P0 to P5, and foot skin samples were collected on P6. The heatmaps and bar graphs, showing the stiffness (kPa) in WT and cKO foot skin at various distances from epidermis. Scale bar, 50 μm. J. Whole-mount immunofluorescent (IF) images of H, showing sweat ducts in green (Krt2) and glands in red (Krt8). Scale bar, 100 µm. K. Quantification of ductal length from J. Each dot represents one duct counted. WT, n = 6 mice; cKO, n = 7 mice. L. IF images of sweat glands in red (Krt8) from H. White dashed lines circle the glandular area. Scale bar, 100 μm. M. Quantification of sweat gland areas from L. Each dot represents one sectioned glandular area. WT, n = 8 mice; cKO, n = 8 mice. Data are presented as mean ± SD. Statistical significance was determined using unpaired two-tailed Student’s t test. *P ≤ 0.05, **P ≤ 0.01, ***P ≤ 0.001.

To identify the structural basis of the stiffness gradient, we used second harmonic generation (SHG) microscopy^28^ to visualize collagen, the predominant ECM component in the skin and a key determinant of tissue stiffness and performed whole-mount imaging of mouse foot skin to map collagen density and its 3D distribution relative to sweat ducts and glands. As shown in Movie S1 and Figures 1D–G, collagen was markedly enriched around the ductal region and much lower near the glands, directly mirroring the spatial pattern of high and low stiffness respectively.

We hypothesized that the stiffness gradient is functionally instructive: high stiffness promotes straight ductal elongation, and low stiffness favors glandular differentiation. To test this, we generated a dermal-specific Collagen1a1(Col1a1) conditional knockout (cKO) mice using Pdgfra^CreER^ strain and confirmed that the dermal stiffness in the Col1a1 cKO dermis decreases overall but still exhibits a gradient pattern by AFM (Figures 1H-I). We performed immunofluorescent (IF) imaging to quantify sweat duct length and sweat gland areas, using keratin 2 (Krt2) and keratin 8 (Krt8) as their markers, respectively. We found that the ductal length is shorter (Figures 1J-K) and the glandular size is bigger (Figures L-M) in Col1a1 cKO footpads. The same pattern was observed in the Col1a1 cKO using a different dermal-specific driver (S100a4^CreER^) (Figures S1E-H), together indicating that the changes in stiffness in the dermal environment significantly impact on glandular shape and size.

### Spatiotemporal transcriptional mapping of mechanical properties and Wnt signaling in dermal fibroblasts across sweat gland morphogenesis

To investigate the temporal dynamics of the mechanical and signaling profiles in the sweat gland microenvironment, we performed single cell RNA sequencing for mouse footpad skin from three critical developmental stages: duct elongation (P0), transition from duct to gland (P3), and glandular maturation (P6) (Figure 2A). We acquired data from a total of 19,125 fibroblasts and annotated their subclusters as papillary dermis (PD), reticular dermis (RD), and hypodermis (HD) according to their respective signature genes (Figures 2B and S2A-C)^27,29^. By analyzing the cell composition at different developmental stage, we found that PD fibroblasts are most dominant during sweat duct elongation (P0) and the percentage of RD fibroblasts increases as glandular differentiation starts after P3 (Figure 2C). The expansion of RD fibroblast at P3 is also supported by high level of proliferation marker gene *Mki67* (Figure S2D). Next, we confirmed their spatial distribution by IF imaging using their respective signature genes: Axin2 for PD^21^, p120 (Ctnnd2) for RD, and Ly6a for HD^29^, confirming their identity and localization within the tissue (Figures 2D and S2C). Notably, we found that the sweat ducts are mostly located in PD and glands are surrounded by RD, whereas the HD lies beneath the glands (Figure 2D). Together, these data suggests that PD and RD may directly influence sweat gland development, but not the HD. From differentially expressed gene (DEG) and KEGG pathway analysis (Figure 2E-F), we identified two outstanding pathways that are elevated in PD: 1) extracellular matrix (ECM)-related pathway, including *Lmna*, *Eln*, *Fn1*, *Col1a1*, and various collagen genes*; 2)* Wnt-signaling pathway, including *Wnt5a*, *Axin2, Rspo3*, and *Lef1* (Figure S2E-F). To further examine the potential correlation of these two pathways, we constructed ECM module using the signature genes and ranked all PD and RD fibroblasts (n = 13,980) based on their ECM module intensity from high to low. The expression levels of *Wnt5a* and *Axin2* for each fibroblast are also mapped to the same plot. We found that PD fibroblasts are enriched at higher ECM level, which progressively decreases in RD fibroblasts. In addition, the expression levels of *Wnt5a* and *Axin2* exhibit a positive correlation with ECM module intensity, also in a gradient pattern, suggesting a mechanistic link between these two pathways (Figure 2G). This spatial gradient is supported by fluorescent in situ hybridization (FISH) for *Wnt5a,* along with the quantification of the fluorescent signal intensity of each positive pixel relative to the distant from the epidermis. We found that *Wnt5a* expression is highest in the PD closest to the epidermis and progressively declined with increasing distance, and higher level at P3 than at P6, indicating the presence of a spatial and temporal gradient (Figure 2H-I).

**Figure 2.**
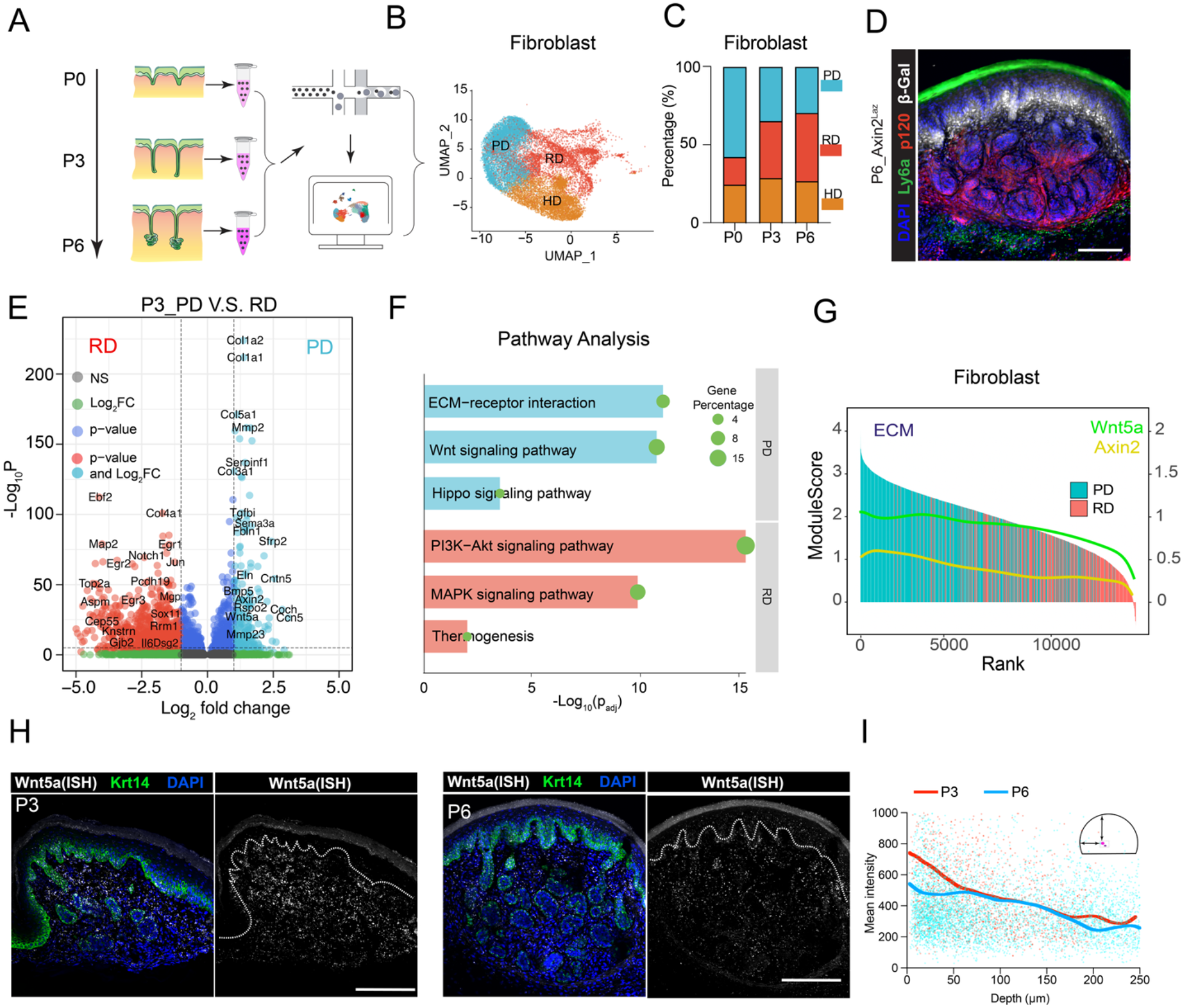
Differential expression of genes involved in mechanical properties and Wnt signaling pathway between papillary and reticular dermis. A. Schematic summary of the experimental workflow. Mouse footpad skin samples containing developing sweat ducts and glands were collected at three critical time points (P0, P3 and P6), followed by tissue digestion and scRNA-seq for transcriptomic analyses. B. UMAP plot of 19,125 fibroblasts, grouped into three unsupervised clusters, representing the papillary dermis (PD), reticular dermis (RD), and hypodermis (HD). C. Stacked bar graph, showing percentage of fibroblasts in PD, RD, and HD clusters across three developmental time points (P0, P3, and P6). D. IF imaging, showing distinct dermal layers in the foot skin using Axin2-LacZ reporter mice. β-galatosidase (white), indicating Axin2 expression and labeling PD; p120 (red) for RD; Ly6a (green) for HD. Scale bar, 50 μm. E. Volcano plot, showing DEGs between PD and RD fibroblasts from P3 mouse foot skin, with absolute fold changes >2 and P <0.05. F. Bar graph, showing significant enrichment of KEGG pathway. The size of each green dot represents the percentage of DEGs associated with the respective KEGG pathway. G. PD and RD fibroblasts (n=13,980) are ranked and aligned according to their ECM module scores. Fitted lines showing *Wnt5a* (green) and *Axin2* (yellow) expression in each fibroblast. The ECM module is composed of *Col1a1, Col3a1, Col5a1, Eln, Vim, Fbn1, Lmna, Fbln1, Fn1,* and *Fbln2*. H. P3(left) and P6(right) foot skin samples stained with DAPI, Krt14, and fluorescent in situ hybridization (FISH) for *Wnt5a*. Dashed lines indicate basement membrane. I. Quantification of *Wnt5a* intensity relative to depth away from epidermis. Red dots indicate data points from P3 (n = 6 mice) foot skin samples, and blue dots from P6 (n = 6 mice). Red and blue lines indicated average mean intensity. Scale bar, 50 μm.

In addition, we found that both ECM and Wnt module level is highest at P0, lower at P3, and lowest at P6, consistent with our finding that ECM and Wnt module levels decline as glandular development progresses to later stage (Figure S2G). Taken together, our findings reveal spatiotemporal changes of mechanical property and Wnt signaling in the glandular tissue microenvironment, suggesting that as sweat bud progenitors invaginate deeper into the dermis at the later stage, they encounter a progressive softer dermal environment and reduced Wnt signaling during morphogenesis.

### High stiffness in the dermis facilitates ductal elongation through upregulation of dermal Wnt5a

To test whether tissue stiffness–dependent morphogenesis represents a conserved mechanism across glandular systems, we cultured epidermal tissues containing mammary, lacrimal, and sweat buds on polyacrylamide hydrogel substrates with varying stiffness. (Figure 3A-G). The stiffness of whole tissue is altered in response to that of the substrates, as measured by AFM, and still exhibit a gradient pattern at the intermediate stiffness levels (Figure 3A-B). We found that the ductal length increases significantly when the tissues are placed on a stiffer substrate, which is consistent across all glands tested (Figure 3A-G).

**Figure 3.**
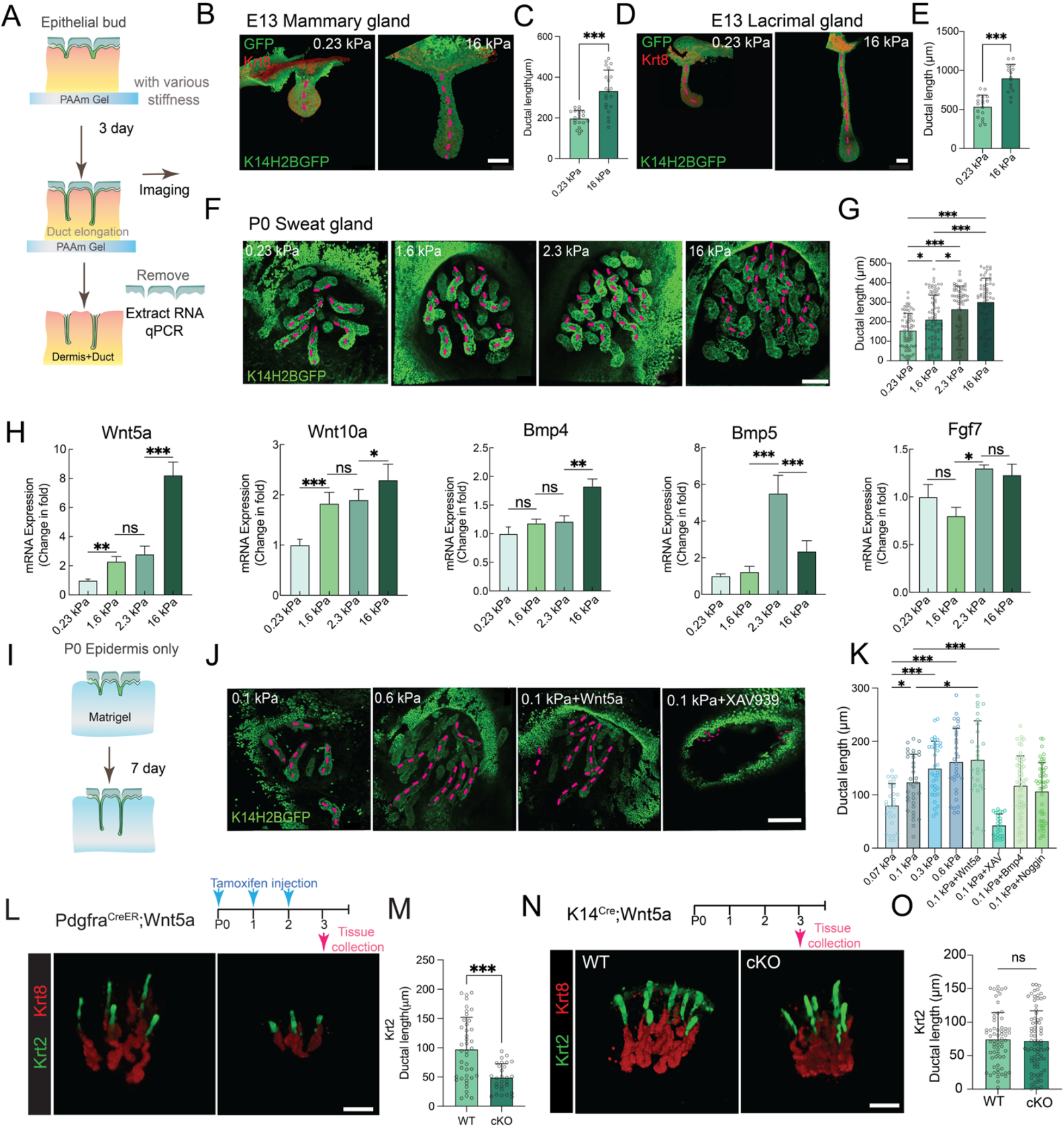
High stiffness promotes sweat duct elongation through upregulation of dermal Wnt5a. A. Schematic summary of the explant culture experimental workflow. B-G. Whole-mount IF images (B, D, F) showing explant cultured on substrates of various stiffness as indicated during the ductal development stages. K14H2BGFP mice were used to label all keratinocytes in green and Krt8 IF staining shown in red. Pink dashed lines indicate the midline of the ducts. Scale bar, 100 µm. (C, E, G) quantification of ductal length based on morphology and midline of the ducts. Each dot represents a single duct. Mammary gland on soft: n = 16 mice; Mammary gland on stiff: n = 18 mice. Lacrimal gland on soft: n = 12 mice; Lacrimal gland on stiff: n = 10 mice. Sweat gland on soft: n = 3 mice, Sweat gland on stiff: n = 3 mice. H. Quantitative PCR analysis of *Wnt5a*, *Wnt10a*, *Bmp4*, *Bmp5* and *Fgf7* expression in the dermal fraction cultured on substrates of varying stiffness. Experimental repeats, n = 3. I. Schematic representation of epidermal sheet (without dermis) explant culture on Matrigel substrates of varying stiffness. J. Whole-mount IF Images of epidermis sheet cultured on varying stiffness Matrigel, with or without Wnt or Wnt inhibitor (XAV939), for 7 days. Pink dashed lines labels the midline of the ducts. Scale bar, 100 µm. K. Quantification of J and Figure S3E. Each dot represents a single duct. n = 3 mice per condition. L. Whole-mount IF images of P3 footpads from Pdgfra^CreER^;Wnt5a^+/+^ (WT) and Pdgfra^CreER^;Wnt5a^fl/fl^ (cKO) mice after 3 days of tamoxifen induction from P0 to P2, showing ducts in green (Krt2) and glands in red (Krt8). Scale bar, 100 µm. M. Quantification of ductal length in L. Each dot represents one duct measured. WT, n = 4 mice; cKO, n = 4 mice. N. Whole-mount IF images of P3 footpads from K14^Cre^;Wnt5a^+/+^ (WT) and K14^Cre^;Wnt5a^fl/fl^ (cKO) mice, showing ducts in green (Krt2) and glands in red (Krt8). Scale bar, 100 µm. O. Quantification of ductal length in N. Each dot represents one duct counted. WT, n = 6 mice; cKO, n = 6 mice. Data are presented as mean ± SD. Statistical significance was determined using an unpaired, two-tailed Student’s t test for (C, E, M, O) and one-way ANOVA for (G, H, K). *P ≤ 0.05, **P ≤ 0.01, ***P ≤ 0.001.

Using scRNAseq data of foot skin containing developing sweat glands, we identified signaling ligands that are expressed in the dermis (PD and RD) (Figure S3C) and examined their expression levels in response to different stiffness in the dermal fractions of tissue explants by quantitative PCR (qPCR). Among these ligands, we found that *Wnt5a, Wnt10a and Bmp4 expression* are elevated with the increasing stiffness, while that of *Bmp2, Bmp5* and *Fgf* ligands do not show correlation with tissue stiffness (Figures 3H and S3D).

Because the dermal fraction of the explants contains both epidermal (sweat bud) progenitors and dermal fibroblasts, we next sought to delineate their respective responses to mechanical and biochemical cues by culturing isolated epidermis containing sweat buds (at P0), excluding the dermis, on Matrigel substrates of varying stiffness with or without specific signaling ligands or inhibitors (Figure 3I). As shown in Figure 3J–K, ductal length increased with rising Matrigel stiffness, although the effect was markedly smaller in magnitude and slower in rate than in full-thickness explants. These findings suggest that while epidermal progenitors retain intrinsic mechanosensing capacity, robust glandular morphogenesis requires the presence of dermal components. To determine whether Wnt and/or Bmp signaling directly regulate ductal elongation in the absence of dermis, we treated epidermal sheet explants with Wnt5a, the Wnt inhibitor XAV939, Bmp4, or the Bmp inhibitor Noggin. Wnt5a treatment markedly promoted ductal extension, whereas XAV939 shortened ducts. In contrast, neither Bmp4 nor Noggin altered ductal growth (Figures 3J-K and S3E), indicating that Wnt signaling plays a pivotal role in driving ductal elongation, whereas Bmp signaling, although active in footpad dermis, appears dispensable in this context.

To examine whether fibroblast-derived Wnt ligands (predominantly Wnt5a, Figure S3C) may promote sweat duct elongation *in vivo*, we generated dermal-specific *Wnt5a* knockout mouse line using Pdgfra^CreER^ and S100a4^CreER^. We first confirmed that the loss of *Wnt5a* in KO dermis, but not in WT dermis, by ISH (Figure S3F). Next, using 3D IF imaging and quantification of ductal length as indicated by Krt2 expression, we found that the ductal length is significantly shorter in *Wnt5a* dermal KO (Figures 3L-M and S3H-I G-H). We further generated K5rtTA; tetO-Dkk1 mice to overexpress Dkk1 for Wnt inhibition, and this similarly resulted in shortened ducts (Figures S3I-J). Although *Wnt5a* is predominantly expressed in dermis, it is also present at a much lower level in the developing sweat duct based on our scRNAseq data (Figure S3C) and ISH image (Figure S3F). To determine whether epidermal (ductal) Wnt5a may contribute to ductal growth, we generated epidermal knockout of *Wnt5a* using K14^Cre^, and confirmed the loss of *Wnt5a* by ISH (Figure S3K). We found that there is no significant difference in the ductal length in *Wnt5a* epidermal KO, compared to their littermate control (Figure 3N-O). These results altogether support the dominant role of dermal Wnt5a in driving ductal elongation upon reception by ductal progenitor cells.

### Low stiffness and Wnt inhibition in the dermis promote glandular formation

As epidermal ducts grow deeper into dermis, progenitor cells at the leading edge progressively encounter a softer microenvironment, coinciding with their transition toward glandular differentiation. To investigate whether mechanical cues modulate this duct-to-gland transition, we performed tissue explant cultures on hydrogels with various stiffness, using developmental stages when ductal elongation was largely complete, and the progenitors were poised for glandular differentiation; E13 for salivary gland and P3 for sweat glands (Figure 4A-G). In contrast to the stiffness-dependent elongation of ductal length observed earlier (Figure 3), increased stiffness now markedly reduced glandular formation, while ductal length remained unchanged across conditions (Figures 4B-D for salivary glands; 4F-G and S4A-B for sweat glands). Furthermore, at E13, a subset of mammary (45%) and lacrimal (22%) glands exhibited early glandular branching, which was significantly inhibited in stiff substrates (Figures S4C-F), supporting the notion that soft mechanical environment facilitate the initiation of glandular morphogenesis. As shown previously in Figure 3H that explant tissues on soft substrate express low level of *Wnt5a*, we found that dermal *Wnt5a* KO mice developed significantly enlarged glands, consistent with explant experiments, with no significant change in ductal length (Figures 4H-K and S4G-J). This change of glandular size is not due to the low expression of *Wnt5a* in the ductal progenitor, as there is no significant difference in glandular size nor ductal length in K14^Cre^;Wnt5a epidermal cKO (Figures S4K-N).

**Figure 4.**
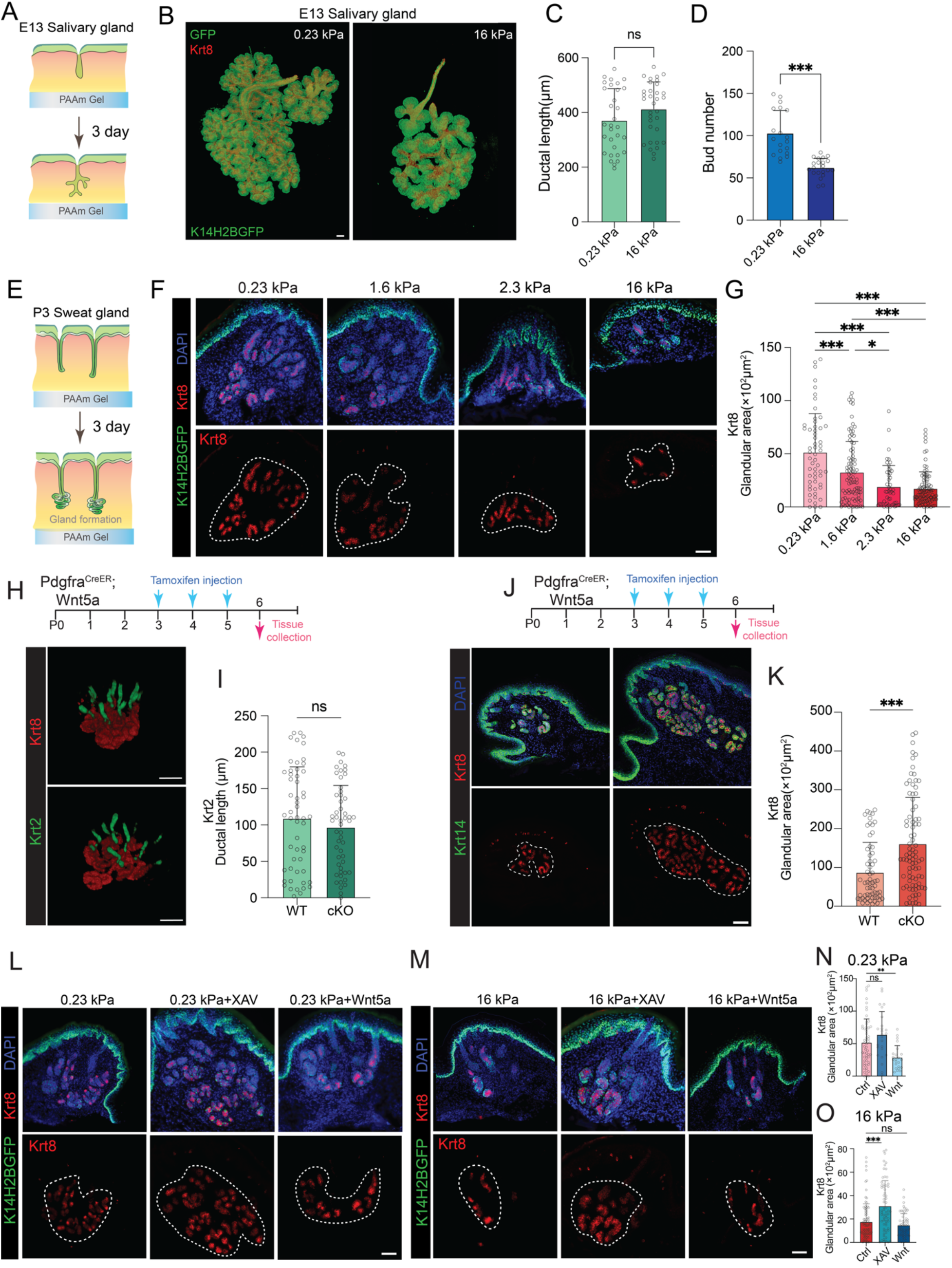
Low stiffness and Wnt inhibition promote glandular formation. A, E. Schematic illustration of the explant culture assays, using E13 salivary gland and P3 foot skin for sweat gland, to examine glandular development. B. Whole-mount IF imaging, showing salivary glands cultured on substrates of different stiffness for 3 days. K14H2BGFP mice were used to label all keratinocytes in green and Krt8 IF staining shown in red. Scale bar, 100 μm. C-D. Quantification of ductal length and bud (acinar) numbers for b. Each dot represents one duct counted. Salivary glands on soft substrate, n = 19 mice; stiff substrate, n = 21 mice. F. IF images, showing Krt8 staining in P3 foot skin explant cultures on substrates of different stiffness for 3 days. White dashed lines circle glandular areas. n = 3 mice per condition. Scale bar, 50 µm. G. Quantification of glandular size in F. Each dot represents one glandular area measured. H-K. Pdgfra^CreER^; Wnt5a^fl/fl^ foot skin, with (cKO) or without (WT) tamoxifen induction during glandular development stage (P3-P6). H. Whole-mount IF images, showing sweat ducts (Krt2) in green and glands (Krt8) in red. Scale bar, 100 µm. I. Quantification of ductal length (Krt2+) in H. Each dot represents one duct measured. WT, n = 5 mice; cKO, n = 4 mice. J. IF images, showing Krt8+ glandular areas circled in white dashed lines. Scale bar, 50 µm. K. Quantification of glandular areas in J. WT, n = 4 mice; cKO, n = 4 mice. Each dot represents one glandular area measured. L-M. IF images, showing P3 foot skin explants, as illustrated in E, cultured on soft (0.23 kPa) or stiff (16 kPa) substrates with or without Wnt inhibitor (XAV) or Wnt5a. White dashed lines circle glandular areas. Scale bar, 50 µm. N-O. Quantification of glandular size by Krt8+ areas in L and M, respectively. Each dot represents one glandular area measured. n = 3 mice per condition. Data are presented as mean ± SD. Statistical significance was determined using an unpaired, two-tailed Student’s t test for (C, D, I, K) and one-way ANOVA for (G, N, O). *P ≤ 0.05, **P ≤ 0.01, ***P ≤ 0.001.

Because both tissue stiffness and Wnt signaling influence glandular size and increased stiffness elevates Wnt5a expression, we hypothesize that Wnt signaling may be downstream of stiffness in directing glandular morphogenesis. To test this hypothesis, we performed the explant experiments using P3 foot skin (as Figure 4E) with a combination of high/low stiffness and Wnt ligand/inhibitor. As shown in Figure 4L-O, consistent with our earlier findings, soft substrates (0.23 kPa) promoted glandular growth; under this condition, Wnt inhibition with XAV939 further enlarged the glands, whereas Wnt5a treatment reduced their size. Conversely, stiff substrates (16 kPa) suppressed glandular growth, Wnt inhibition restored gland size, while Wnt5a addition had no significant effect. Collectively, these findings demonstrate that reducing both tissue stiffness and Wnt signaling synergistically promotes glandular growth, and that the impact of Wnt signaling on glandular progenitors can override the effects of tissue mechanical cues.

### Dermal Piezo1 regulates Wnt5a expression and thereby supports ductal fate

To investigate the mechanisms by which dermal fibroblasts sense stiffness gradients in the niche and respond by expression of *Wnt5a*, we sought to identify the key factor that mediates this process. First, we examined the expression of all mechanosensing-related ion channels in fibroblast clusters using our scRNAseq data and found that Piezo1 is expressed at the highest level among them (Figure S5A). We confirmed that the increase of stiffness upregulates Piezo1 expression in both skin tissue explant and fibroblast cell culture (Figures S5B-D). Pharmalogical modulation further showed that Wnt5a expression in fibroblasts increases with the Piezo1 agonist Yoda1 and decreases with the antagonist GsMTx4 (Figure S5E), indicating that Wnt5a levels in dermal fibroblasts are regulated by Piezo1 activity.

Next, to determine whether defects in dermal mechanosensing affects ductal development *in vivo*, we generated a conditional knockout Piezo1 using Pdgfra^CreER^ and found that cKO exhibit significantly shorter ducts compared to controls at P3 (Figures 5A-B). Consistently, dermal tissues from Piezo1 cKO mice showed a blunted mechanosensitive response, with reduced *Wnt5a* and *Axin2* expression on both soft and stiff substrates relative to WT littermates. This deficit was even more pronounced under stiff conditions (Figures 5C-D).

**Figure 5.**
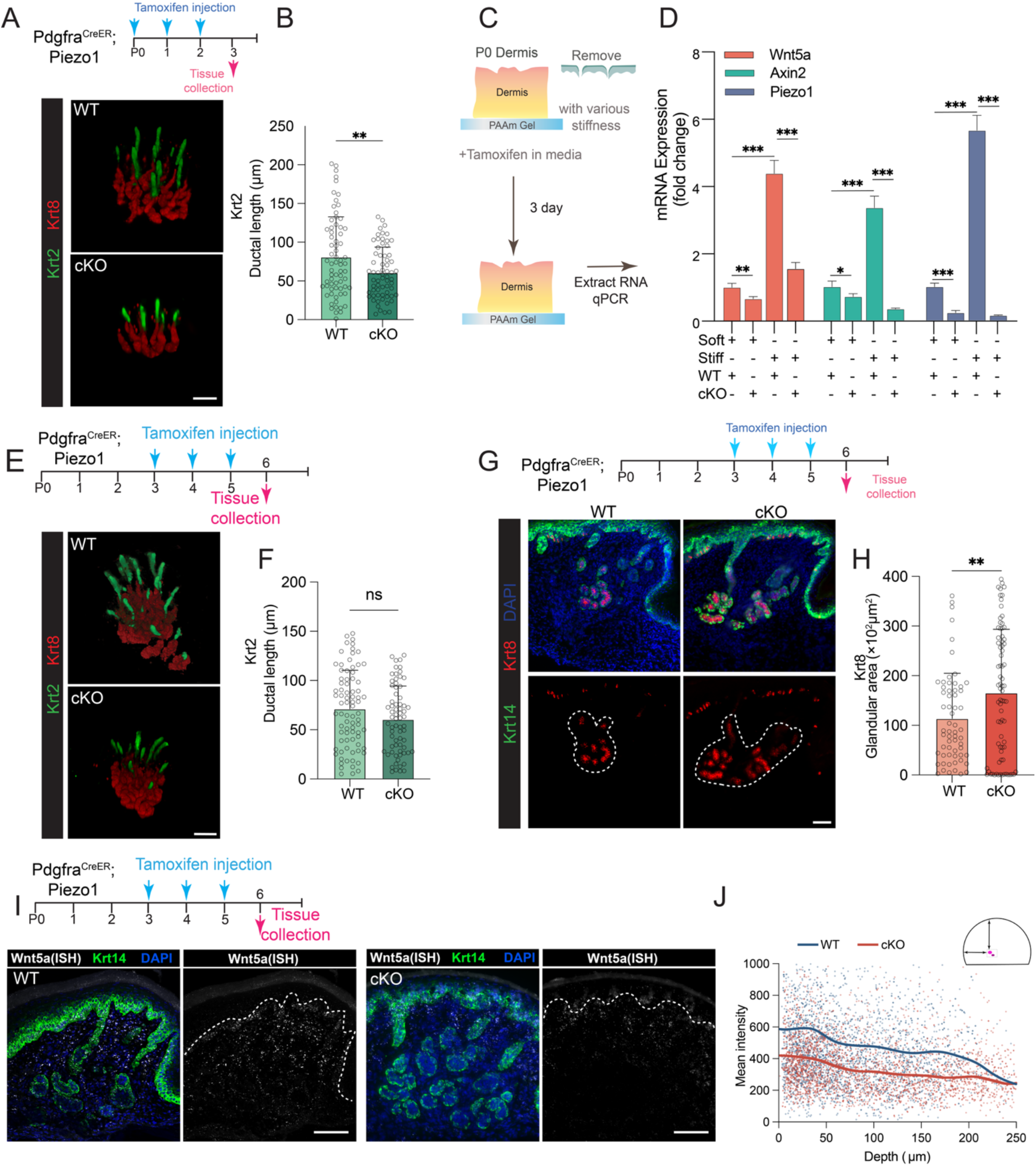
Dermal fibroblasts convert stiffness gradient into Wnt5a gradient via Piezo1 mechanosensing. A. Whole-mount IF images, showing sweat ducts and glands from P3 foot skin of WT and Pdgfra^CreER^; Piezo1^fl/fl^ , after 3 days of tamoxifen induction from P0-P2. Scale bar, 100 µm. B. Quantification of ductal length using Krt2 in green. Each dot represents one duct measured. WT, n = 5 mice; cKO, n = 4 mice. C. Schematic of dermal explants experiment using WT and Pdgfra^CreER^; Piezo1^fl/fl^ mouse foot skin tissues on stiff or soft substrates. D. Bar graph, showing expression of Wnt5a, Axin2 and Piezo1 expression in cultured dermal explants under the indicated conditions by qPCR analysis. Experimental repeats, n = 3. E-J. P6 foot skin of WT and Pdgfra^CreER^; Piezo1^fl/fl^ , after 3 days of tamoxifen induction from P3 to P5. E) Whole-mount IF images, showing sweat ducts (Krt2) in green and glands (Krt8) in red. Scale bar, 100 µm. F) Quantification of ductal length (Krt2+) in e. Each dot represents one duct measured. WT, n = 6 mice; cKO, n = 4 mice. G) IF images, showing Krt8+ glandular areas circled in white dashed lines. Scale bar, 50 µm. H) Quantification of glandular areas in G. Each dot represents one glandular area measured. WT, n = 5 mice; cKO, n = 5 mice. I) Fluorescent in situ hybridization (FISH) of Wnt5a in white. J) Quantification of the mean intensity of Wnt5a in relation to the distance from epidermis. WT, n = 4 mice; cKO, n = 4 mice. Data are presented as mean ± SD. Statistical significance was determined using an unpaired, two-tailed Student’s t test for (B, F, H) and one-way ANOVA for (d). *P ≤ 0.05, **P ≤ 0.01, ***P ≤ 0.001.

To specifically assess the role of dermal mechanosensing in glandular differentiation, we ablated Piezo1 after P, when ducts are already formed. Under this condition, ductal length was unchanged (Figures 5E-F), whereas glandular size was significantly increased (Figure 5G-H). In situ hybridization further revealed that loss of dermal Piezo1 attenuates the Wnt5a gradient, with overall reduced expression (Figures 5I-J and S5F-G), suggesting that a basal Wnt5a gradient is reinforced by Piezo1-dependent mechanosensing in the dermis.

Together, these data indicate that dermal fibroblasts utilize Piezo1 to sense tissue stiffness and regulate Wnt5a expression, thereby directing glandular progenitor cell fate specification through Wnt signaling.

### Ductal vs glandular cell fate specification operates as a gradient in response to the level of Wnt signaling

Thus far, we demonstrated that tissue stiffness regulates Wnt5a expression in fibroblasts, and together these cues specify ductal versus glandular fate. Because both stiffness and Wnt5a form gradients in the dermis, invaginating progenitors are exposed to a continuum of Wnt signals during morphogenesis. To test whether the duct-to-gland transition reflects a threshold response or a graded continuum and to define the range of effective Wnt concentration, we engineered a 3D microfluidic system to recapitulate the glandular microenvironment *in vitro*. Conventional two-channel, single-layer devices^30^ generate gradients confined to a narrow region (<5 mm), limiting cellular exposure and biasing toward binary outcomes (Figure S6A). While multi-channel designs improve gradient linearity, their structural complexity makes them difficult to reproduce^31^ (Figure S6B). To address these limitations, we developed a microfluidic sweat gland development model (µSgDM) that integrates a multi-channel PDMS gradient generator with a 50 μm micropillar array, simplifying device architecture while enabling robust, linear gradients across a large 3D culture area without compromising structural integrity.

COMSOL was used to simulate the dimensions of the channels and culture areas to reach the most optimal gradient outcome (Figures 6A and S6C-D). The concentration in each channel can be given by:

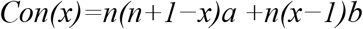

where x is the inlet order number of the cell culture area, n is the number of microfluidic layers, and a and b are the concentrations of the two main inlet molecules. Further, this µSgDM employs an eight-layer sequential diffusive mixing mechanism to establish a stable and linear Wnt signaling gradient across a wide culture field, thereby better mimicking the spatial scale of the native dermal environment (Figure 6B). µSgDM-generated gradient was further validated through trypan blue diffusion testing (Figure 6C), demonstrating enhanced stability and linearity.

**Figure 6.**
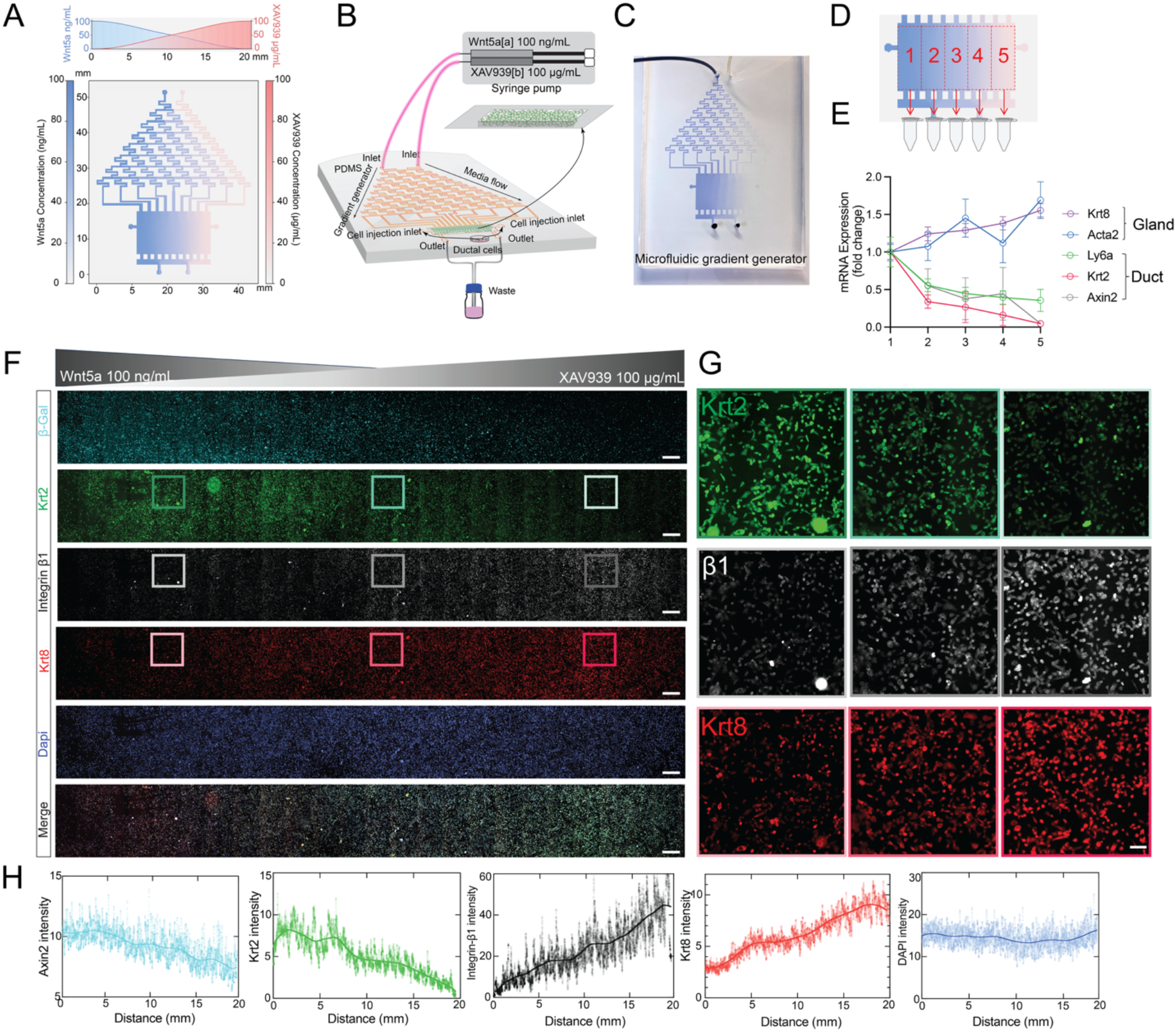
Duct-to-gland cell fate specification in response to Wnt gradient. A. COMSOL simulation, showing distribution of the morphogen gradient. Top: Quantification of Wnt5a and Xav939 concentration along the length of the culture chamber. Bottom: Spatial distribution within the chamber. B. Schematic of the microfluidic device to generate morphogen gradient. E media containing the morphogen gradient flows laminarly from the microfluidic channels into the cell culture chamber containing ductal progenitors embedded in Matrigel. The media inflow is controlled by precise syringe pumps, while used media exits from the opposite side of the cell chamber into a waste container. C. Image of the device and chemical gradient generated by the microfluidic device, visualized by trypan blue. D. Diagram illustrating collection of cells from 5 regions within the culture areas within different levels of Wnt5a and XAV939. E. Expression of ductal (Ly6a, Krt2, Axin2) and glandular (Krt8, Acta2) signature genes within these 5 regions from D were analyzed by qPCR. Experimental repeats, n = 6. F-G. IF images, showing expression of ductal and glandular genes in the culture area of the gradient generator. Boxed areas are shown in higher magnification in G. Scale bars, 500 µm for F and 100 µm for G. H. Quantification of fluorescent intensity of indicated markers across the culture area.

Purified ductal progenitor cells were embedded in Matrigel and seeded onto the culture area of the µSgDM and subjected to a gradient of *Wnt5a* (0–100 ng/mL) and the Wnt inhibitor XAV939 (0– 100 µg/mL) from the other direction. The culture area was divided into five regions (sample 1–5), and gene expression within each was analyzed by qPCR (Figure 6D). After 72 hours of culture in the gradient device with decreasing Wnt5a concentration and increasing XAV939 concentration, we observed decreased expressions of ductal marker genes (Axin2, Ly6a, and Krt2), as well as increased expressions of glandular marker genes (Krt8 and Acta2) by qPCR (Figure 6E).

Next, using IF imaging across the entire culture area (∼22 mm), we first validated the Wnt gradient with ductal progenitor cells from Axin2-LacZ reporter mice, which exhibited progressively reduced reporter activity with decreasing *Wnt5a* and increasing Wnt inhibitor. Consistent with our qPCR results, the expression of ductal marker (Krt2) gradually declined, while that of glandular markers (Krt8 and Integrin β1) increased along the gradient (Figure 6F-H). Together, these findings demonstrate the dominant role of Wnt signaling in directing sweat ductal v.s. glandular fate specification in a graded manner.

## Discussion

Although morphogen gradients are fundamental to nearly all aspects of development and organogenesis, replicating true gradients in experimental systems has proven highly challenging. Most previous studies have relied on a limited set of incremental morphogen concentrations or mathematical modeling to approximate these gradients^11,13^. Existing microfluidic platforms for gradient generation remain constrained by design limitations: highly intricate architectures can produce precise gradients but often compromise reproducibility, whereas simpler devices lack the ability to establish stable and linear gradients across a broad concentration range^30,31^. To better recapitulate the dynamic morphogen gradients encountered by progenitor cells during organogenesis, and to enable both computational simulations and experimental validation, we engineered an efficient eight-layer microfluidic device integrating micropillars to provide a uniform culture space and prevent collapse within large chambers, while maintaining low fabrication complexity. This design supports the generation of stable and continuous morphogen gradients, ensures uniform cell distribution and robust mechanical support, and importantly, facilitates systematic investigation of morphogen signaling responses at both cellular and tissue scales.

The critical role of mechanotransduction in regulating epithelial morphogenesis has been demonstrated in several developmental contexts, such as intestinal crypt formation^32^, lung branching^33^, hair placode formation^23^. However, these insights have largely been derived from cultured cell assays or organoid system, which remain markedly different from the complexity of *in vivo* developing tissues and organs. To bridge this long-standing gap, we performed tissue-macroscale mechanical profiling using AFM and 3D ECM imaging, enabling direct visualization and quantification of mesenchymal mechanical properties – an aspect often overlooked in cell-based or organoid models. Notably, we identified for the first time that a gradient of tissue stiffness exists in the mesenchyme as a prerequisite for establishing a morphogen gradient. This stiffness gradient is spatially pre-patterned prior to epidermal mini-organ initiation (Figure S1) and exhibits temporal dynamics during maturation (Figure 2). The resulting mesenchymal morphogen gradient specifies epidermal cell fate in a graded manner, as progenitors proliferate and collectively invaginate deeper into the dermis, and differentiate into lineage-specific, functionally distinct cell types required for complete organogenesis.

We demonstrated that tissue stiffness in the skin microenvironment is primarily sensed by dermal fibroblasts, which act as critical translator of ECM architecture and mechanical property into biochemical cues to control epidermal cell fate. Our work added to a long-standing paradigm that dermal fibroblasts impact on epidermal cell fate specification, especially through expressing Wnt5a, whose mechanosensitive regulation appears to be a conserved mechanism across multiple developmental systems and species in patterning and morphogenesis. In vertebrate limb development, Wnt5a is expressed in mesenchymal tissues in a graded manner and is required for proper proximodistal patterning, coordinating epithelial invagination and mesenchymal stiffness^34^. Similar roles have been found in the developing marsupial (sugar glider) flight membranes, where Wnt5a mediates directional cell behaviors and progenitor zone organization in response to local mechanical or structural cue^35^. These findings, together with ours, underscore the context-dependent yet evolutionarily convergent role of Wnt5a as a key mediator that transmits mesenchymal mechanosensing to epithelial cell fate decisions during organogenesis.

Only recently have mechanical cues been appreciated as key regulators of exocrine gland morphogenesis, particularly in the budding and branching of compound tubuloacinar glands. In mammary organoids, matrix stiffness influences cell fate and branch elongation^36,37^, while in the salivary gland, epithelial-basement membrane adhesion dominates over cell-cell interaction during budding^38^. In contrast, the patterning and morphogenesis of simple tubular glands, such as intestinal^39^, gastric glands^39,40^, endometrial^41^, and eccrine sweat glands^27,42^, remain less understood. Despite being the most abundant appendage in human skin and essential for thermoregulation and fluid balance, there has been no evidence for *de novo* sweat gland regeneration in adult skin after injury, making *in vitro* generation a longstanding goal for treating anhidrosis or severe burns. Here, we establish a mechanistic paradigm in which tissue mechanics instruct biochemical signaling to coordinate glandular morphogenesis. We define the coupled mechanical and biochemical requirements that drive the sequential formation of ductal and glandular compartments with distinct functions and identify a general principal of exocrine gland morphogenesis - mesenchymal mechanical cues actively shape epithelial cell fate and organ architecture. These findings provide a conceptual framework for engineering diverse glandular epithelia for therapeutic applications.

## Methods

### Mice

All mouse procedures were performed in accordance with animal protocols approved by Institutional Animal Care and Use Committee (IACUC) at NYU Grossman School of Medicine. The following strains were used: Axin2^tm1^(cre/ERT2)^Rnu^/J (Axin2-Cre^ERT2^)^43^; Gt(ROSA)26Sortm9(CAG-tdTomato)Hze/J (RosaLSL-tdTomato)44; Piezo1tm2.1Apat/J (Piezo1flox)45; Wnt5a^tm1.1Krvl^/J (Wnt5a^flox^)^46^, were purchased from the Jackson laboratory. Tg(Pdgfra-cre/ERT)467Dbe/J (Pdgfra-Cre^ERT2^)^47^; Axin2^tm1Wbm^/J (Axin2^LacZ^)^43^; (S100a4-Cre^ERT2^)^48^; Col1a1^tm1c^(EUCOMM)^Wtsi^/RKlJ (Col1a1^flox^), were kind gifts from Dr. Mayumi Ito (NYU Grossman School of Medicine).

Tg(KRT5-rtTA)T2D6Sgkd/J (Krt5-rtTA)^49^ mice and Hprt1^tm1^(tetO–Dkk1)^Spdl^ (tetO-Dkk1) mice were kind gifts from Dr. Sarah Millar (Mount Sinai).

Tg(KRT14-cre)1Amc/J (K14-Cre)^50^ and Tg(KRT14-HIST1H2BJ/GFP)18Efu (K14-H2BGFP)^51^ were kind gifts from Dr. Elaine Fuchs (Rockefeller University).

To induce CreER activity in pups, tamoxifen (TAM) treatment was performed via intraperitoneal injection (i.p.) (50 mg kg^-1^body weight) of a 10 mg ml^-1^ solution in corn oil per day. For tetracycline-inducible mouse models, mice were given doxycycline-containing chow (20g/kg) (Bio-Serv, Flemington, NJ) for the indicated time period.

### Explant culture

Polyacrylamide hydrogels with varying stiffness were prepared as previously described in For tetracycline-inducible mouse models, mice were given doxycycline-containing chow (20g/kg) (Bio-Serv, Flemington, NJ) for the indicated time period. glass-bottom dishes^52,53^. Fibronectin was conjugated onto the gel surface by immersing the hydrogels in 1 mL solution of 0.5 mg/mL Sulfo-SANPAH (Pierce) in PBS, followed by activation under UV light for 10 minutes. After activation, the gels were washed with PBS and incubated in 50 μg/ml fibronectin (Sigma) for 1 hour at room temperature.

Mouse foot skin tissues were dissected from P0 and P3 mice, gently placed dermal-side down on different stiffness substrates, and cultured in the explant culture media along with pharmacological agents, including Wnt5a (R&D, 50 ng/mL), XAV939 (Sigma, 5 µg/mL) Bmp4 (R&D, 50 ng/mL), Noggin (R&D, 25 µg/mL), Calyculin A (Sigma, 25 nM), Blebbistatin (Abcam, 25 µM) if applicable^54^. Foot skin tissues were cultured on hydrogel substrates at 37°C for 72 hours before being embedded in OCT blocks or fixed in 4% paraformaldehyde (PFA) in PBS. All culture experiments were performed at least three times with three skin samples (each from a different embryo) per condition.

### AFM measurements

We employed an AFM (Oxford Instruments), mounted on the stage of an inverted microscope (Zeiss AxioObserver Z1), to assess freshly excised foot skin tissue. The AFM cantilever, with a spring constant of 0.09 N/m and a 5 μm diameter glass sphere attached to its tip (Novascan), was navigated over the tissue with the aid of light microscopy. Controlled deformations were applied, and compressive forces were measured via cantilever deflection. The elastic modulus (E) of the tissue was determined by fitting the contact region of the force curves to a Hertz model for a spherical indenter of radius. For each skin sample, 18 × 18 points were probed within a 100 × 100 μm region, with at least three regions analyzed per sample, in both the papillary and reticular dermis. To assess whether AFM probing affected the tissue’s elastic properties, repeated indentations were performed at single points in preliminary tests. Spatial variations in dermal stiffness were visualized using color-coded force maps.

### RNA extraction and qPCR analysis

After culture, explant skin tissues were incubated in 50 mM EDTA for 20 min at 37°C, and epidermis and dermis were physically separated then snap-frozen in liquid nitrogen. The frozen epidermis and dermis were crushed with tissue manual pulverizer respectively, RNA was then extracted with a RNeasy Mini Kit (Qiagen), which quantified with spectrophotometer (Nanodrop 8000). cDNA was synthesized from the reverse transcribed mRNA using SuperScript VILO cDNA Synthesis Kit (Invitrogen), followed by real-time PCR that mixing cDNA with SYBR Green PCR Master Mix (Applied Biosystems) and primers, performing on the ABI Prism 7900 HT sequence detection system (Applied Biosystems). Hprt1 was used as a housekeeping gene for normalization, and the results were analyzed using the 2^−ΔΔCt^ method. Primer sequences for qPCR were listed in Table1.

### Immunofluorescence staining

Embedded tissues were cryosectioned into 10 μm, and fixed in 4% PFA for 10 min, washed in PBS, incubated in blocking buffer (1% gelatin, 1% bovine serum albumin, 2.5% normal donkey serum, 2.5% normal goat serum, 0.3% Triton in PBS) for 1 hour at room temperature (RT). The sections were incubated with primary antibodies overnight at 4°C. After 3 times wash with PBS, samples were stained with the Alexa Fluor 488/555/647-conjugated (1:500, Invitrogen) secondary antibodies for 1 hour at RT, and nuclei were stained with DAPI (1:10000, Molecular Probes) for 5 min. Samples were mounted in Prolong Gold Antifade Mount to preserve the fluorescent signals from quenching (Invitrogen). Slides were examined on microscope (Zeiss Axio Imager Z2) equipped with an argon–krypton laser, and images were collected using a 20 × objective.

To measure the duct length in different culture conditions, the whole foot skin tissues were fixed in 4% PFA overnight at 4°C and follow the CUBIC tissue clearing protocol that described before^55^. Briefly, after delipidation by CUBIC-1 reagent, staining with Krt2 and Krt8 primary antibodies was performed for three days at RT following another three days of incubation with corresponding secondary antibodies. Last, refractive index (RI) matching with CUBIC-2 reagent (RI: 1.55) was introduced to have cleared footpads. Whole mount fluorescent images were acquired with a confocal microscope (Zeiss LSM 880), 20× objective was used to collect images. To detect the collagen deposition in the footpads, cleared K14H2BGFP footpads were imaged by Leica Stellaris Confocal microscope equipped with MaiTai DeepSee laser for second harmonic generation (SHG). Collagen within 10 μm of ductal and glandular edges was classified as ductal collagen and glandular collagen, respectively. The total area for each category was quantified from individual frames.

The following primary antibodies were used: Krt14 (Chicken, 1:800, BioLegend), Krt2 (Rabbit, 1:200, Abcam), Krt8 (Rat, 1:500, DSHB), Ly6a (Rat, 1:100, Invitrogen), p120 (Mouse, 1:100, Invitrogen), β-Gal (Chicken, 1:800, Abcam), Integrin β1(Hamster, 1:100, Biolegend), Piezo1 (Mouse,1:200, Novus).

### Fluorescent In situ hybridization (FISH) and quantification of Wnt5a expression gradient

To detect Wnt5a RNA expression, an antisense RNAscope probe targeting mouse Wnt5a (ACDBio) was introduced to section samples. We used the Multiplex Fluorescent reagent Kit (ACDBio) with TSA Vivid Fluorophore 520 (ACDBio) to amplify the *Wnt5a* signals. To be compatible with general IF staining, we applied RNA-Protein Co-Detection Ancillary Kit (ACDBio) to co-stain Krt14 as a reference of keratinocytes. All procedures were executed following the manufacturer’s manuals. The images were taken by the confocal microscope (Zeiss LSM 880) with z step of 2 µm. To quantify the *Wnt5a* expression gradient, we first used Krt14 channel as a mask of basal layer. In addition, DAPI channel was referenced as cells or tissues.

Then, a square (32 × 32 pixel) was introduced to scan through the whole image to take measurement of mean intensity of *Wnt5a* channel upon the region-of-interest was situated in the mesenchyme/outside of the basal layer. A border line of epidermis was then generated manually to calculate the closest distance of the region-of-interest to the epidermis. Last, *Wnt5a* expression (detected by RNAscope probe) and the corresponding distance to the epidermis were then plotted to visualize the gradient within mesenchyme of mouse food pads.

### Preparation of single cell suspensions

Foot skin samples from K14H2BGFP mice (P0, P3, and P6) were dissected and the epidermis and dermis, including sweat glands, were separated by incubating the skin in 50 mM EDTA at 37 °C for 20 minutes. The epidermal and dermal tissues underwent distinct digestion procedures. The epidermis was digested with 0.25% Trypsin/EDTA for 10 minutes, washed in 4% FBS/PBS, filtered through a 40 µm cell strainer, and centrifuged at 300 g for 10 minutes at 4 °C. The dermal tissue was digested in 2 mg/mL collagenase/HBSS (Hanks’ Balanced Salt Solution) at 37 °C with gentle agitation on a horizontal shaker for 1 hour. After removing the supernatant by centrifugation, the tissue was further digested with 0.25% Trypsin/EDTA for 10 minutes at 37 °C. The dermal suspension was filtered through a 40 µm nylon strainer, neutralized with 4% FBS/PBS, and centrifuged under the same conditions as the epidermis. The resulting cell suspensions were prepared for fluorescence-activated cell sorting (FACS) to enrich for GFP-positive cells.

### Single-cell library construction and sequencing

The sorted cellular suspensions were submitted to the Genome Technology core facility at NYU Langone and loaded on a 10x Genomics Chromium instrument to generate single-cell gel beads in emulsion (GEMs). Approximately 10K cells were loaded per channel. Single-cell RNA-Seq libraries were prepared using the following Single Cell 3’ Reagent Kits v3.1: Chromium Next GEM Single Cell 3’ GEM, Library & Gel Bead Kit v3.1, PN-1000121; Chromium Next GEM Chip G Single Cell Kit, PN-1000120 and Single Index Kit T Set A PN-1000213 (10x Genomics) and following the Single Cell 3’ Reagent Kits v3.1 User Guide (Manual Part # CG000204 Rev D). Once the cDNA libraries were generated, they were subject to Illumina NovaSeq 6000 paired-end sequencing.

### Single-cell RNA-seq data processing and analysis

The transcriptomes of 12046, 19982, and 13946 live single cells from P0, P3, and P6 of single-cell RNA-seq data were processed by Cell Ranger Single Cell Software Suite (version 1.3), barcode ad UMI processing, and single-cell 3′ gene counting.

UMI for each cell was analyzed using the Seurat package (version 5.0.1) in R (version 4.2.0). Cells with fewer than 200 detected genes or more than 10% mitochondrial gene expression were excluded from further analysis. The resulting gene-cell matrices were log-normalized using Seurat’s NormalizeData function. After normalization, variable genes were identified using the FindVariableFeatures function, with selection based on variance stabilizing transformation (vst) of gene expression. Data were scaled using ScaleData, and principal component analysis (PCA) was performed to reduce dimensionality with the top 2,000 most variable genes. Significant principal components (PCs) were determined using an elbow plot, and the first 10 PCs were selected for downstream analysis. Cells were clustered using a graph-based clustering algorithm (FindClusters function) with the resolution parameter set to 0.5, followed by visualization using the Uniform Manifold Approximation and Projection (UMAP) via the RunUMAP function. Cell-type annotation was performed by cross-referencing canonical marker genes with published datasets and validated using the FindMarkers function for differential expression testing.

Differentially expressed genes were identified using the Wilcoxon rank-sum test, with an adjusted P < 0.05 considered significant. For integration of multiple datasets, the FindTransferAnchors and MapQuery functions were employed using P3 as the reference, allowing for combined analysis across different time points. To assess the biological relevance of identified clusters, functional enrichment analysis was performed using Kyoto Encyclopedia of Genes and Genomes (KEGG) pathway analysis. The module score was evaluated by the AddModuleScore function.

### Microfluidic Channel Fabrication

SU-2000 SERIES negative photoresist (MICROCHEM) was used to create a mold for the microfluidic channel. SU-2010 was spin-coated onto a cleaned large glass slide (NC1824406, Ted Pella Inc.) at 3000 r.p.m. for 60 s with a 300 r.p.m./s ramp, producing a 10 µm thick layer. The film was soft-baked at 65°C for 2 min and then at 95°C for 3 min. The Heidelberg uMLA Maskless Aligner exposed SU-8 at 800 mJ/cm² using a 365 nm LED. The exposed sample underwent a Post Exposure Bake (PEB) at 95°C for 4 min, then cooled before being developed in SU-8 Developer (Kayaku) for 90 s. The developed sample was rinsed with 2-propanol, dried with N₂ gas, and hard-baked at 150°C to relieve residual stress. The patterned sample was salinized with Trichloro(1H,1H,2H,2H-perfluorooctyl) silane (448931, Sigma Aldrich) in a desiccator overnight. The PDMS (Sylgard 184, Dow Corning) microfluidic channel was prepared by mixing PDMS and curing agent in a 10:1 ratio, casting on the patterned SU-8 glass slide, and curing at 65°C for 4 h to transfer the SU-8 pattern to PDMS. The inlets and outlets of the microfluidic channel were punched using Standard Biopsy Punches (3331AA, Integra). The cured PDMS and cleaned large cover glass (NC0719784, Ted Pella Inc.) were exposed to oxygen plasma for 40 s under conditions of 30 W, 30 mL/min oxygen flow, and 800 mTorr chamber pressure to enhance surface bonding. The treated glass and PDMS bonded easily. A leak test was conducted by flowing DI water through both inlets of the microfluidic channel at 1 mL/min by using syringe pump (1000-US, New Era Pump Systems Inc.), confirming no leakage.

### Flow simulation

The finite-element method (FEM) platform COMSOL Multiphysics (version 6.1) was used to model the microfluidic channel. The simulation incorporated the Laminar Flow and Transport of Diluted Species modules. A DXF file of the real-size microfluidic channel geometry, created in AutoCAD, was imported, with channel height set to 10 µm. The designed microfluidic channel includes two inlets and two outlets to generate a stable concentration gradient in the main chamber, with two additional side openings for cell seeding. The main chamber is supported by 50 µm diameter pillars spaced 1 mm apart to prevent roof collapse. A physics-controlled fine mesh was applied, and the simulation was performed using a steady-state model. Material properties were set to those of water, with the diffusion coefficient of diluted particles at 10^-10^ m²/s to match experimental conditions. Wall boundary conditions were set to no-slip, with both inlets having an inflow velocity of 10^-5^ m/s one inlet with a diluted particle concentration of 1 mol/m³ and the other at 0 mol/m³. Outlets were configured to suppress backflow. The simulation was computed using the MUMPS solver.

### Ductal cell culture assay

P2 neonates from Axin2-Laz mice were sacrificed, and the foot skin was harvest. The digestion method of dermis with the sweat ducts is the same as Single cell suspension method. The cells were resuspended and prepared for FACS using Sca1 (BD Biosciences, 1:100) and CD49f (Invitrogen, 1:200) markers to identify ductal cells. After sorting, the ductal cells were cultured on mitomycin C-treated J2 feeder cells in 0.05 mM low calcium media.

### Quantification and statistical analysis

ImageJ (NIH) was used to investigate ductal length and glandular area. Statistical analyses were performed with Prism software (GraphPad 10.2.3) or R studio (version 4.2.0) is used for coding. Statistical significance was evaluated with the two-tailed unpaired Student’s t-test for comparing two groups, and one-way ANOVA for comparing more than two groups. *P ≤ 0.05, **P ≤ 0.01, ***P ≤ 0.001; ns, not significantly different.

## Acknowledgements

We thank Dr. Chen W. at NYU Tandon School of Engineering for initial discussions; NYU Genome Technology Center for sequencing and raw data processing (P30CA016087); NYU Microscopy Laboratory for assistance with imaging (S10RR023704); MSKCC Molecular Cytology for AFM measurements; Dr. Lim C and Dr. Ma C for helpful discussions and technical support.

## Funding

NYU Hansjӧrg Wyss Department of Plastic Surgery start-up grant (C.L.). NIH NIAMS of R01AR080136 and R01AR082033E (C.L.). Mallinckrodt Jr. Foundation, I. T. Hirschl Trust (C.L.). NIGMS of R35GM147406 (H.C).

## Author Contributions

J.T., C.L. conceived the project, designed the experiments and wrote the manuscript. M.L. performed SHM imaging and *in situ* hybridization, generated all the mouse strains and collected all mouse samples. M.D. assisted the mouse experiments. M.X., S.M and M.I. provided valuable mouse strains. N.D. and H.C. designed and made the microfluidic device. J.T performed the rest of experiments and all analyses and simulations. C.L supervised the project and secured funding. All authors provided insightful input and have read and approved the final manuscript for publication.

## Competing interests

The authors declare no competing interests.

## Data and materials availability

Data availability Current data can be found in Gene Expression Omnibus (NCBI GEO, www.ncbi.nlm.nih.gov/geo) GSE279851. All data acquired and generated by the lab are recorded in the manuscript and the supplementary materials, or available from the corresponding author upon reasonable request.

## Supplementary Materials

**Figure S1.**
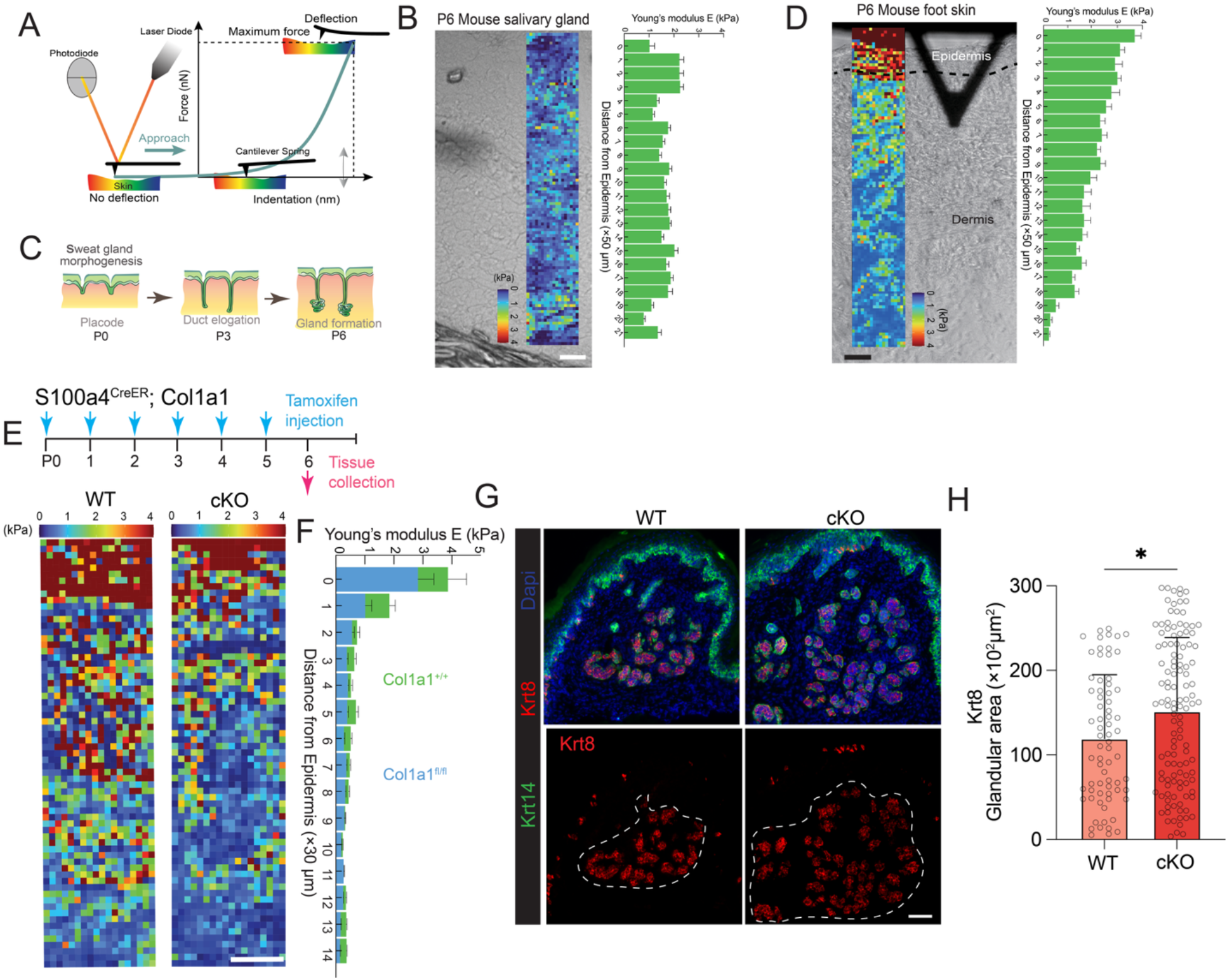
Dermal stiffness controls the glandular morphology. A. The scheme of AFM, illustrating stiffness measurement of tissue cross section (10 μm thickness). The AFM cantilever was navigated over specific region of the tissue. Force-displacement curves were generated at each point by plotting cantilever deflection over controlled deformation. B. Distribution of Young’s modulus (E) in P6 mouse salivary gland (glandular area), visualized as color-coded heat maps and bar graphs, showing the quantified values in kPa and aligned according to their corresponding distances from epidermis. n = 3 mice. Scale bar, 50 µm. C. Diagram, illustrating critical stages during sweat gland morphogenesis at P0 (placode), P3 (ductal elongation) and P6 (glandular differentiation). D. Distribution of Young’s modulus (E) in P6 mouse foot skin (area with no sweat glands), shown in color-coded maps with corresponding quantification data. n = 4 mice. Scale bar, 50 µm. E-H. P6 foot skin samples from Col1a1 WT and cKO mice, with tamoxifen induction from P0 to P5. E) Heatmap showing the distribution of stiffness. Scale bar, 50 µm. F) corresponding quantification data. WT, n = 4 mice; cKO, n = 5 mice. G) IF images, showing Krt8 staining in red to indicate the glands. White dashed line circles the glandular area. Scale bar, 50 µm. H) Quantification of Krt8+ glandular area in g. Each dot represents one glandular area measured. WT, n = 6 mice; cKO, n = 6 mice. Data are presented as mean ± SD. Student’s t test, unpaired, two tailed was used to determine the statistical significance. *P ≤ 0.05, **P ≤ 0.01, ***P ≤ 0.001.

**Figure S2.**
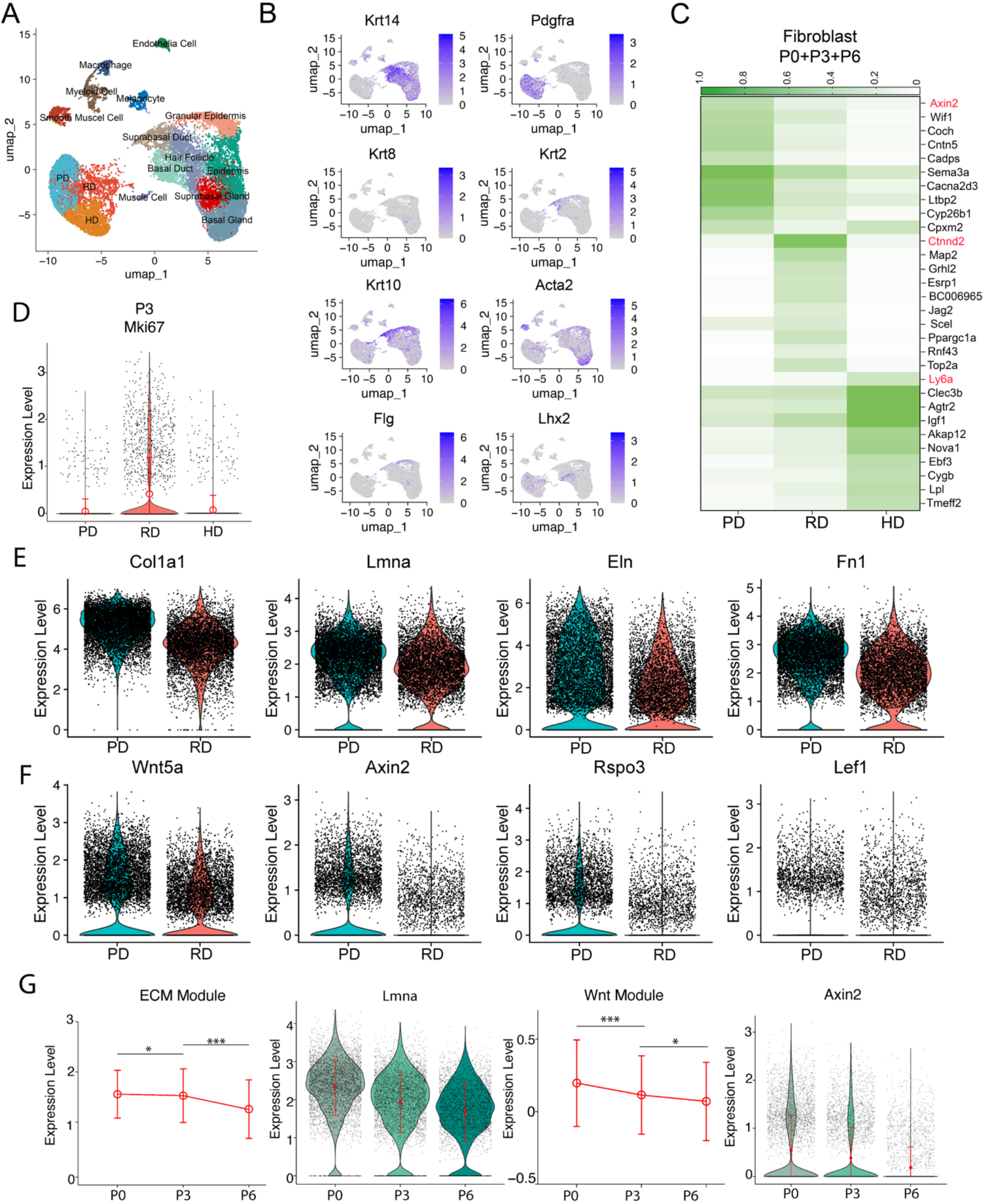
Dermal fibroblast gene expression dynamics across different regions and developmental timepoints. a. UMAP plot of P3 single cell RNA-seq data, grouped into unsupervised clusters, color-coded by cell type. b. Feature plots, showing expression of signature genes in different clusters for annotation. c. Heatmap showing the top 10 differentially expressed genes (DEGs) in PD, RD, and HD clusters. Genes highlighted in red were validated by IF in Figure 2D. d. Violin plot, showing expression of Mki67 in PD, RD, and HD populations at P3, indicating fibroblasts in RD are actively proliferating. E. Violin plots, showing expression of key genes in extracellular matrix (ECM) modules in PD and RD. F. Violin plots, showing expression of key genes in Wnt signaling modules in PD and RD. G. Violin plots, showing the temporal changes of ECM and Wnt signaling gene modules at critical developmental time points P0, P3, and P6. Wnt module encompassed *Wnt5a, Fzd1, Dvl1, Rspo2, Rspo3, Axin2, Dvl2, Ctnnb1, Lef1,* and *Tcf4*.

**Figure S3.**
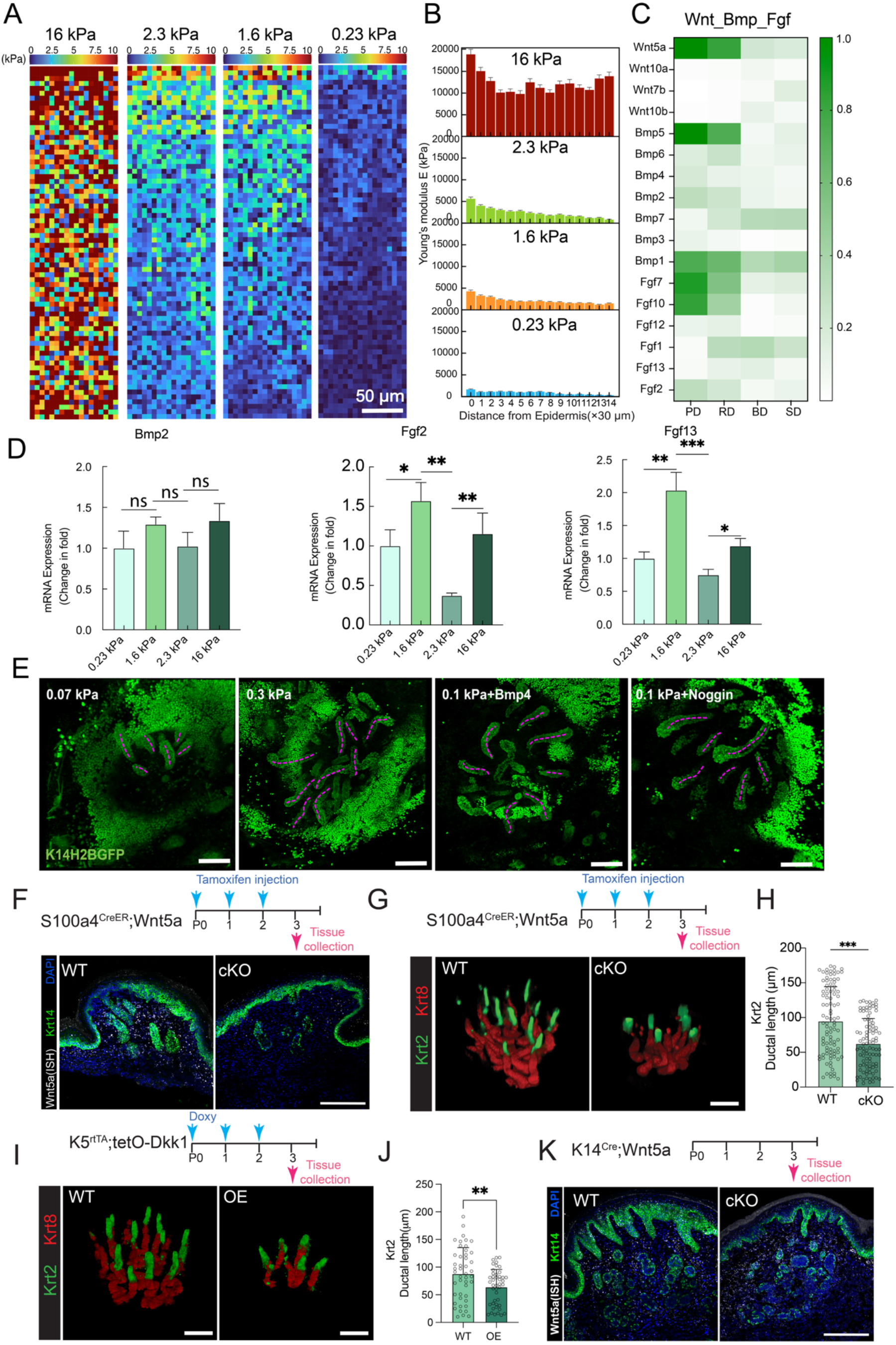
High stiffness promotes sweat duct elongation through elevating dermal Wnt5a. A. Spatial distribution of Young’s modulus (E), showing the tissue stiffness of the P0 foot skin after culturing for 3 days on substrates of different stiffness as indicated in color-coded maps. Scale bar, 50 μm. Note that tissue stiffness still exhibits a spatial gradient in the medium range (2.3 and1.6 kPa). B. Quantification of foot skin stiffness in A, showing stiffness in areas away from epidermis. n = 3 mice per condition. C. Heatmap showing expression of Wnt, Bmp and Fgf ligands in dermis (PD and RD) and sweat ducts (BD, Basal Ductal cells; SD, Suprabasal Ductal cells). D. Bar graphs, showing Bmp2, Fgf2, and Fgf13 expression in cultured explants under the indicated conditions by qPCR. Data are presented as mean ± SD. Statistical significance was determined using one-way ANOVA. *P ≤ 0.05, **P ≤ 0.01, ***P ≤ 0.001. Experimental repeats, n = 3. E. Whole-mount IF images of epidermis sheet cultured on Matrigel of different stiffness, treated with or without Bmp4 or its inhibitor (Noggin) for 7 days. Pink dashed lines label the midlines of the ducts which were used for quantification. Scale bar, 100 μm. F. ISH and IF images, showing expression of Wnt5a in P3 foot skin from S100a4CreER;Wnt5a+/+ (WT) and S100a4CreER;Wnt5afl/fl (cKO) mice. Scale bar, 50 µm. G. Whole-mount IF images of P3 footpads from S100a4CreER;Wnt5a+/+ (WT) and S100a4CreER;Wnt5afl/fl (cKO) mice after 3 days of tamoxifen induction from P0 to P2, showing ducts in green (Krt2) and glands in red (Krt8). Scale bar, 100 µm. H. Quantification of ductal length in G. Each dot represents one duct measured. WT, n = 7 mice; cKO, n = 5 mice. I. Whole-mount IF images of P3 footpads from K5rtTA;tetO (WT) and K5rtTA;tetO-Dkk1 (OE) mice after 3 days of doxy induction from P0 to P2, showing ducts in green (Krt2) and glands in red (Krt8). Scale bar, 100 µm. J. Quantification of ductal length in I. Each dot represents one duct measured. WT, n = 3 mice; cKO, n = 3 mice. K. ISH and IF images, showing expression of Wnt5a in P3 foot skin from K14Cre;Wnt5a fl/fl mice. 996 Scale bar, 50 µm.

**Figure S4.**
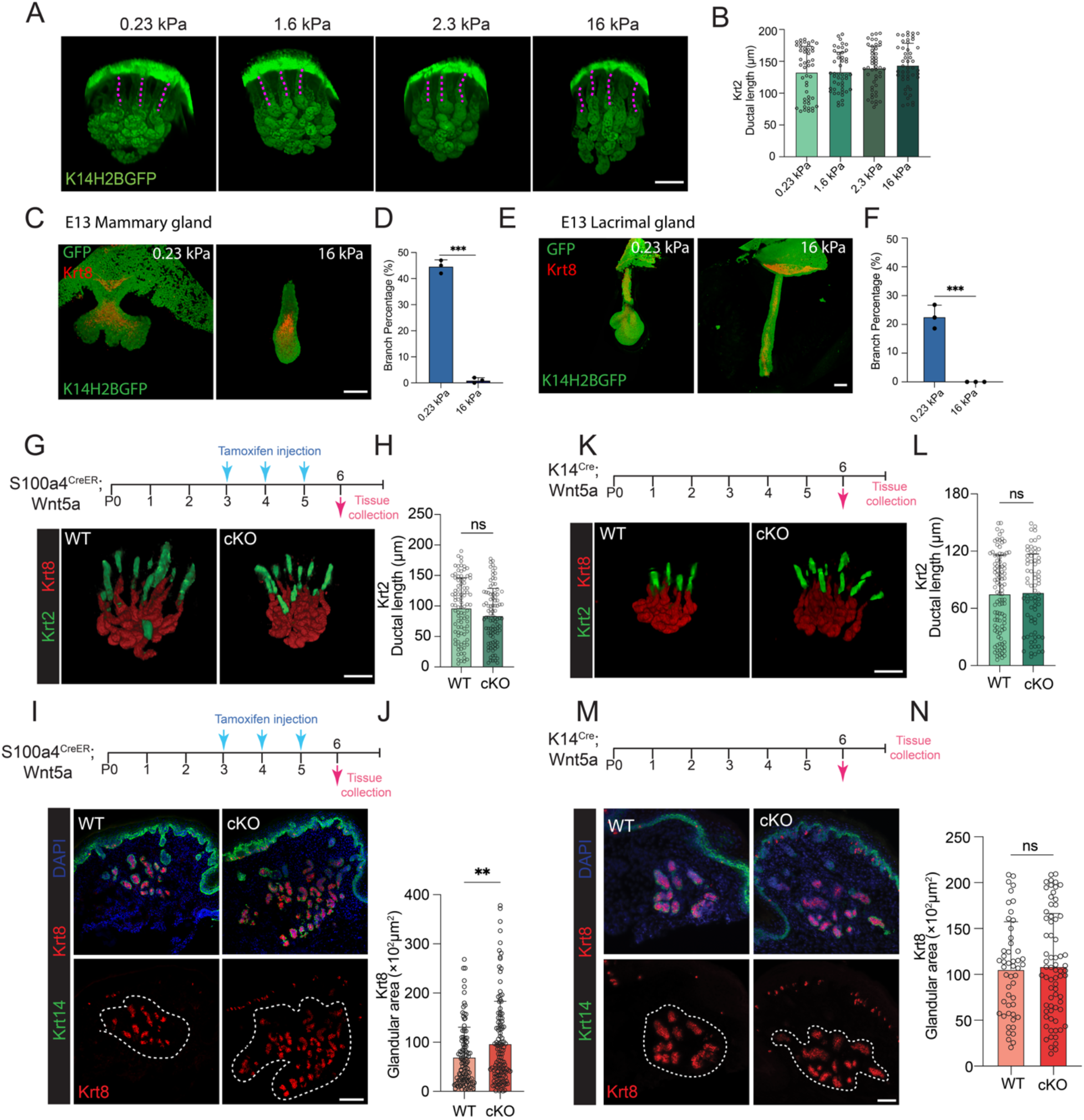
Low stiffness and decreased dermal Wnt signaling promote glandular formation. A. Whole-mount IF images showing the P3 foot skin explant cultured on substrates of varying stiffness as indicated for 3 days. K14H2BGFP mice were used to label all keratinocytes in green. Pink dashed lines indicate the midline of the ducts. Scale bar, 100 µm. B. Quantification of a. Each dot represents a single duct. n = 3 mice per condition. C-F. Whole-mount IF images showing (C, E) E13 mammary and lacrimal gland explant cultured on different stiffness substrates for 3 days. K14H2BGFP mice were used to label all keratinocytes in green and Krt8 IF staining shown in red. (D, F) Quantification of branch percentage based on morphology. Mammary gland on soft: n = 16 mice; Mammary gland on stiff: n = 18 mice. Lacrimal gland on soft: n = 12 mice; Lacrimal gland on stiff: n = 10 mice. Scale bar, 100 μm. G-J. P6 foot skin of WT and cKO (S100a4CreER; Wnt5afl/fl), tamoxifen induction during glandular development stage (P3-P6).(G) Whole-mount IF images, showing sweat ducts (Krt2) in green and glands (Krt8) in red. Scale bar, 100 µm. (H) Quantification of ductal length (Krt2+) in K. Each dot represents one duct measured. WT, n = 6 mice; cKO, n = 6 mice. (I) IF images, showing Krt8+ glandular areas circled in white dashed lines. Scale bar, 50 µm. (J) Quantification of glandular areas in I. WT, n = 8 mice; cKO, n = 7 mice. Each dot represents one glandular area measured. K-N. P6 foot skin of WT and cKO (K14Cre; Wnt5afl/fl). (K) Whole-mount IF images, showing sweat ducts (Krt2) in green and glands (Krt8) in red. Scale bar, 100 µm. (L) Quantification of ductal length (Krt2+) in K. Each dot represents one duct measured. WT, n = 6 mice; cKO, n = 6 mice. (M) IF images, showing Krt8+ glandular areas circled in white dashed lines. Scale bar, 50 µm. (N) Quantification of glandular areas in M. Each dot represents one glandular area measured. WT, n = 7 mice; cKO, n = 6 mice. Data are presented as mean ± SD. Statistical significance was determined using unpaired two tailed Student’s t test. *P ≤ 0.05, **P ≤ 0.01, ***P ≤ 0.001

**Figure S5.**
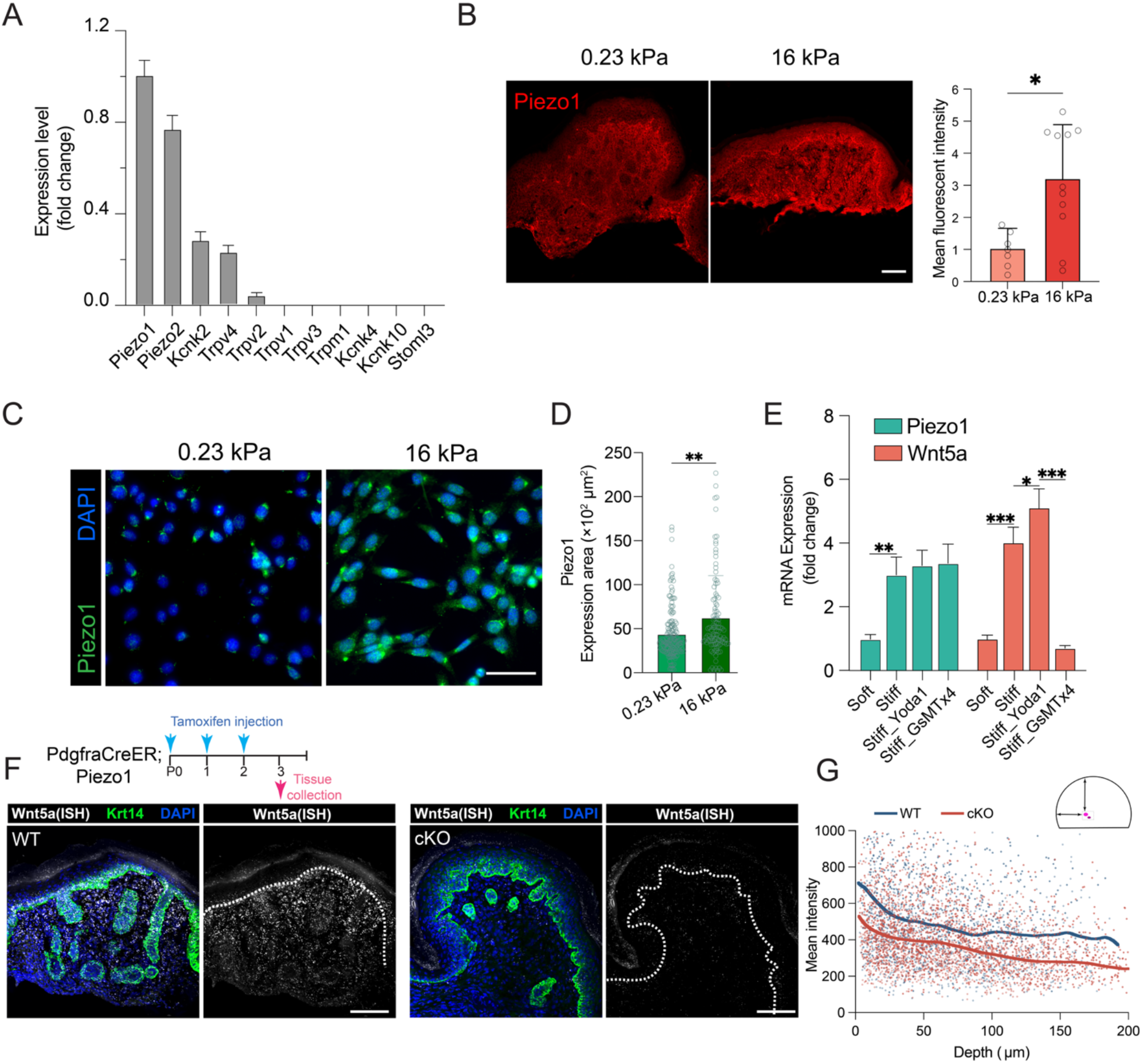
Piezo1-mediated mechanosensing regulates Wnt5a expression in dermis. A. Single cell RNA sequencing (scRNA-seq) analysis, showing expression levels of mechanosensitive ion channel genes in fold change in dermal fibroblasts. Data are presented as mean ± SEM. B. IF images, showing Piezo1 expression in P3 foot skin explants cultured on soft (0.23 kPa) and stiff (16 kPa) substrates for 3 days, and normalized mean intensity quantification data on the right, WT, n = 3 mice; cKO, n = 3 mice. Scale bar, 50 μm C-D. IF images (C) and quantification (D) of Piezo1 expression in dermal fibroblasts cultured on soft and stiff substrates for 3 days. Experimental repeats, n = 4. Scale bar, 50 μm. E. qPCR analysis of Piezo1 and Wnt5a in dermal fibroblasts on soft and still substrates, with or without Yoda1 (Piezo1 agonist) and GsMTx4 (Piezo1 antagonist). F-G. P6 foot skin of WT and cKO (PdgfraCreER;Piezo1fl/fl) foot skin samples stained with DAPI, Krt14, and FISH for Wnt5a. (F) FISH and IF images, showing Wnt5a gradient in WT, which is diminished in cKO. (G). Blue dots indicate data points from WT samples, and the blue line indicates the fitted regression line (n = 4 mice); red dots indicate data points from cKO samples, and the red line indicates the fitted regression line (n = 7 mice). Scale bar, 50 μm. Data are presented as mean ± SD. Statistical significance was determined using an unpaired,two-tailed Student’s t test for (D) and one-way ANOVA for (E). *P ≤ 0.05, **P ≤ 0.01, ***P ≤ 0.001

**Figure S6.**
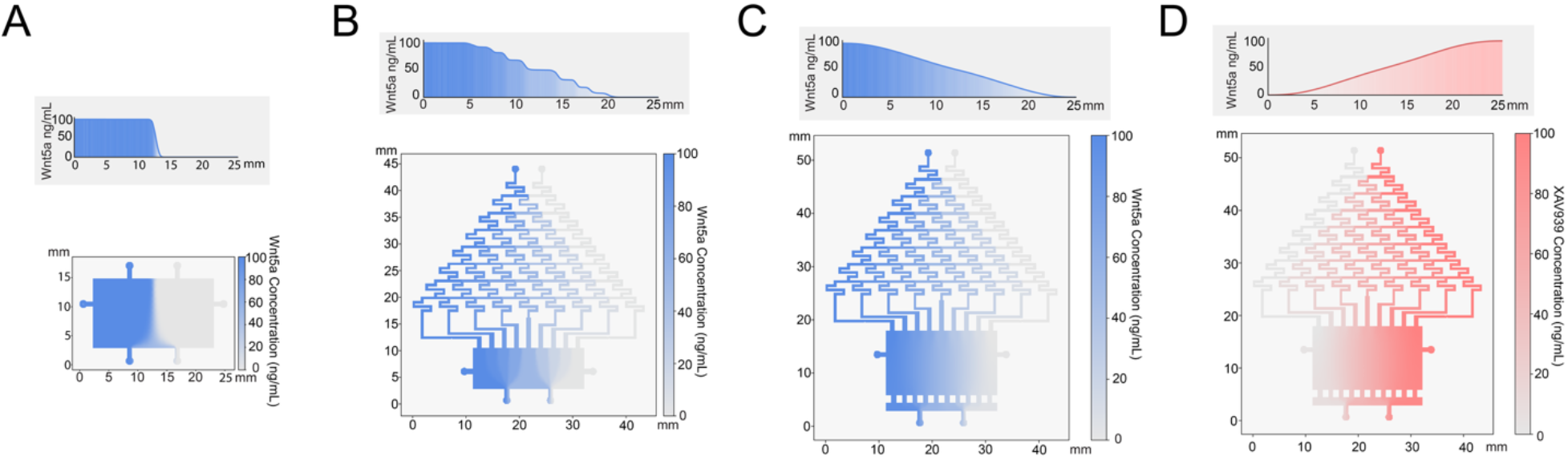
µSgDM generates a linear gradient in simulations. A–D. COMSOL simulations, showing morphogen gradient profiles in different gradient generating devices. (A,B) conventional devices. (C,D) µSgDM. Top: Quantification of morphogen concentration along the length of the culture chamber. Bottom: device designs and morphogen concentration within the chamber.

**Table 1.**
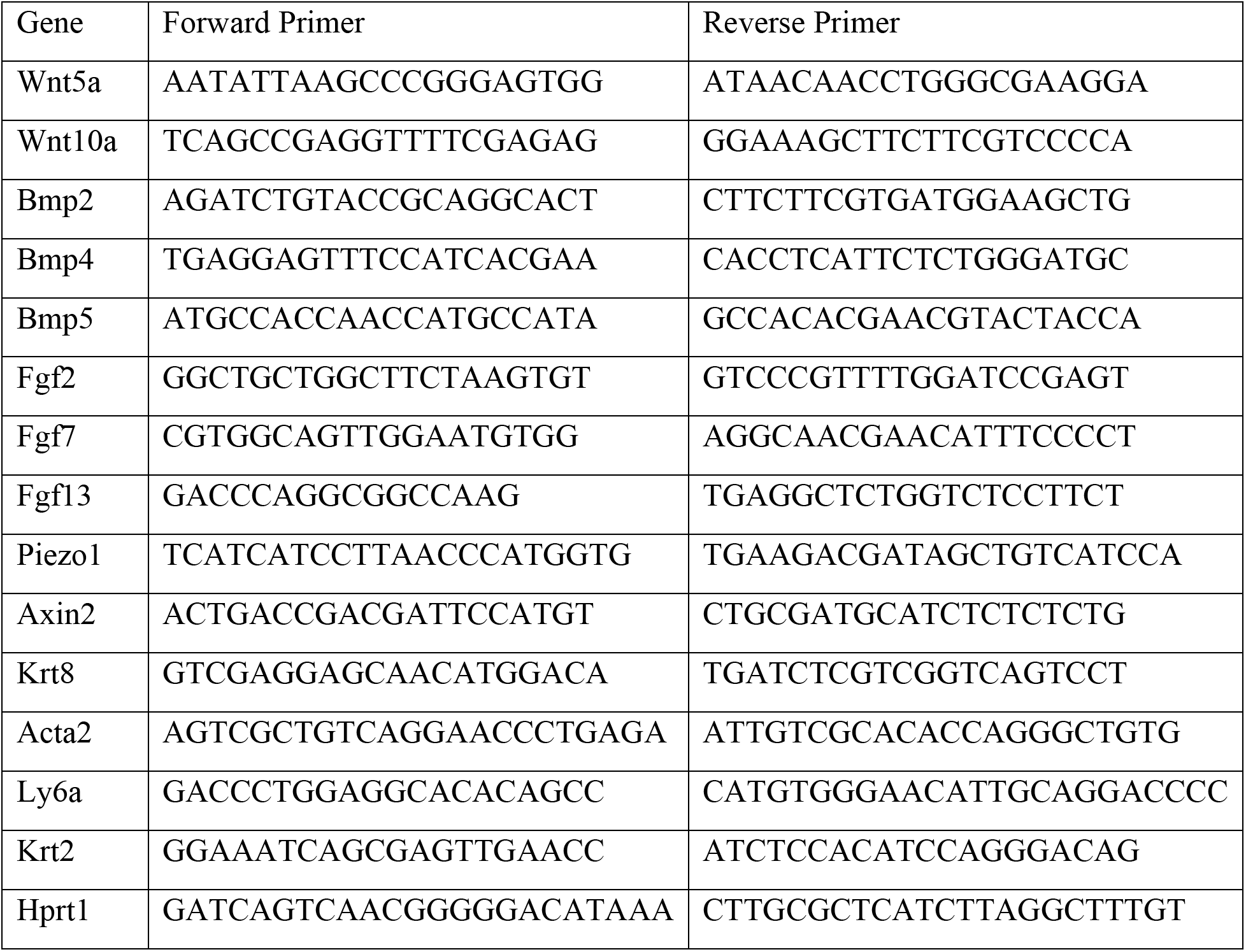
Primer sequences for qPCR analysis.

## Notes

### Competing Interest Statement

The authors have declared no competing interest.

